# Endogenous Rab38 regulates LRRK2’s membrane recruitment and substrate Rab phosphorylation in melanocytes

**DOI:** 10.1101/2022.06.20.496629

**Authors:** Alexandra Unapanta, Farbod Shavarebi, Jacob Porath, Carson Balen, Albert Nguyen, Josh Tseng, Yiyi Shen, Michelle Liu, Pawel Lis, Santiago M. Di Pietro, Annie Hiniker

## Abstract

Point mutations in LRRK2 cause Parkinson’s Disease and augment LRRK2’s kinase activity. However, cellular pathways that enhance LRRK2 kinase function have not been identified. While overexpressed Rab29 draws LRRK2 to Golgi membranes to increase LRRK2 kinase activity, there is little evidence that endogenous Rab29 performs this function under physiological conditions. Here we identify Rab38 as a novel physiological regulator of LRRK2. In mouse melanocytes, which express high levels of Rab38, Rab32, and Rab29, knockdown of Rab38 but not Rab32 or Rab29 decreases phosphorylation of multiple LRRK2 substrates, including Rab10 and Rab12, by both exogenous and endogenous LRRK2. In B16-F10 mouse melanoma cells, Rab38 drives LRRK2 membrane association, and overexpressed kinase-active but not kinase-inactive LRRK2 shows striking pericentriolar recruitment, which is dependent on the presence of endogenous Rab38 but not Rab32 or Rab29. Deletion or mutation of LRRK2 at the Rab38 binding site in the N-terminal armadillo domain decreases LRRK2 membrane association, pericentriolar recruitment, and ability to phosphorylate Rab10. Consistently, overexpression of LRRK2_350-550_, a fragment that encompasses the Rab38 binding site, blocks endogenous LRRK2’s phosphorylation of Thr73-Rab10. Finally, disruption of BLOC-3, the guanine nucleotide exchange factor for Rab38 and 32, inhibits Rab38’s regulation of LRRK2. In sum, our data identify Rab38 as a physiologic regulator of LRRK2 function and lend support to a model in which LRRK2 plays a central role in Rab GTPase coordination of vesicular trafficking.

## Introduction

Leucine-rich repeat kinase 2 (LRRK2) mutations are one of the most common genetic drivers of autosomal dominant Parkinson’s Disease (PD), causing ∼5% of familial PD.^1^ LRRK2 encodes a 286 kDa protein with two catalytic domains: a ROC GTPase domain linked by a COR domain to a serine-threonine kinase.^2^ The remainder of the LRRK2 protein consists of protein-protein interaction domains (armadillo [ARM], ankyrin [ANK], LRR, and WD40 repeats) as well as a regulatory loop region between the ANK and LRR domains that can dictate LRRK2 binding partners and subcellular localization.^3^ How LRRK2 drives PD is not known. However, important work suggests, first, that LRRK2 kinase activity is increased by PD-driving mutations, and second, that 14 Rab proteins (including Rab3, 5, 8, 10, 12, 29, 35, and 43) are LRRK2 kinase substrates and share a conserved Ser/Thr phosphorylation site.^4, 5^ Consistently, hyperactivation of LRRK2 by PD-driving mutations causes defective ciliogenesis and centrosomal cohesion, both of which are Rab-mediated processes.^6–9^ Rab GTPases are critical modulators of intracellular vesicular and endolysosomal trafficking, suggesting that LRRK2’s misregulation of endolysosomal pathways may be involved in its disease-driving function.^10^

In addition to being a LRRK2 substrate, overexpressed Rab29 (also called Rab7L1), increases LRRK2 kinase activity in cellular models.^11^ This is consistent with evidence from animal models showing that Rab29, which falls in the PD risk locus PARK16, genetically interacts with LRRK2 to cause neurodegeneration.^12, 13^ Further, Rab29 protein and LRRK2 physically interact, with Rab29 directly binding LRRK2’s ARM domain, particularly LRRK2_360-450_.^14–16^ Very recent work also identified the extreme N-terminus of LRRK2 as a high-affinity binding site for LRRK2-phosphorylated Rabs, suggesting a feed-forward pathway causing LRRK2 and phospho-Rab accumulation at membranes.^15^ In the cell, overexpressed Rab29 drives overexpressed LRRK2 kinase activation by increasing membrane localization of LRRK2.^11^ A mitochondrially targeted Rab29 construct also activates LRRK2, supporting that Rab29-mediated LRRK2 activation is independent of precise membrane identity.^17^ Critically, almost all studies thus far demonstrate that knockdown/knockout of endogenous Rab29 in multiple cell lines does not decrease LRRK2 kinase function and does not augment Parkinsonian phenotypes in animals, calling into question the physiologic role of Rab29 in LRRK2 activation.^18, 19^ An important exception is a study of Rab29 in the mouse macrophage cell line Raw264.7, in which Rab29 knockdown decreases LRRK2 activation following lysosomotropic stress, suggesting that Rab regulation of LRRK2 may be cell-type and context dependent.^20, 21^

Rab32 and Rab38 are highly homologous to Rab29 (56% and 52% identity to Rab29, respectively). Like Rab29, both bind to LRRK2’s ARM domain in vitro.^14^ Rab32 also co-immunoprecipitates with LRRK2 when co-expressed in cells.^22^ In vitro, LRRK2 complex formation with Rab32/Rab38 requires GTP-bound Rab, suggesting that LRRK2 might be an effector of these small GTPases.^14^ Despite this possibility, the functional relationship between Rab32/Rab38 and LRRK2 has not been thoroughly investigated. Rab32 and Rab38 lack the conserved Ser/Thr LRRK2 phosphorylation site present on Rab29 and there is no evidence they are LRRK2 substrates.^4^ The only assessment of endogenous Rab32/Rab38’s ability to regulate LRRK2 was performed in mouse embryonic fibroblasts (MEFs).^19^ In this work, LRRK2 kinase activity after combined Rab32/Rab29 knockdown/knockout was not different from LRRK2 kinase activity in Rab29 knockout or WT MEFs.^19^ However, MEFs express no Rab38 and low levels of Rab32.^19^ Thus, the effect of Rab38 on LRRK2 activity was not tested at all and the system might not have been optimal for measuring the effect of Rab32.

We postulated a role for Rab32 and/or Rab38 in regulating LRRK2 function for a number of reasons. First, Rab32 and Rab38 directly bind to LRRK2’s ARM domain in vitro^14^ and Rab32 co-immunoprecipitates with LRRK2 in cells.^22^ Second, Rab32 and Rab38 have restricted tissue expression and are highly expressed in cells that produce lysosome-related organelles (LROs), such as melanocytes (producing melanosomes), alveolar Type II pneumocytes (producing lamellar bodies), and numerous inflammatory cells including macrophages and neutrophils.^23, 24^ All of these cell types additionally express high levels of LRRK2, potentially suggesting a functional interaction.^3, 25–27^ Most importantly, LRRK2 knockout or LRRK2 kinase inhibition causes striking buildup of enlarged lamellar bodies in alveolar Type II pneumocytes in animal models, including non-human primates.^28, 29^ LRRK2 kinase inhibition appears to precisely phenocopy the effect of Rab38 mutation/knockout (cht mouse, Ruby rat), and both show enlarged and abundant lamellar bodies.^30–32^ This genetic interaction strongly implicates LRRK2 and Rab38 in a common pathway related to biogenesis of at least some lysosome-related organelles (LROs).

Here we demonstrate that Rab38 but not Rab32 or Rab29 endogenously regulates LRRK2’s kinase function and subcellular localization in murine melanocytes. In these cells, knockdown of Rab38 but not Rab32 or Rab29 decreases both overexpressed and endogenous LRRK2’s phosphorylation of Rab substrates. Rab38 but not Rab32 or Rab29 drives pericentriolar recruitment of exogenous kinase-active LRRK2 in B16 melanoma cells. Knockdown of Rab38 but not Rab32 or Rab29 inhibits recruitment of kinase-active LRRK2 in B16 cells to pericentriolar membranes, which in turn decreases pericentriolar recruitment of endogenous Rab32 and phosphorylation of endogenous Rab10. Consistently, inhibition of Rab38’s effects on LRRK2—either by disrupting the LRRK2-Rab38 interaction through LRRK2 domain/point mutations or by loss-of-function mutation/knockdown of BLOC-3, a guanine nucleotide exchange factor for Rab38/Rab32—recapitulates these effects. Our data thus identify Rab38 as an important upstream Rab GTPase that regulates LRRK2 under physiologic conditions. Furthermore, they add support to a model in which LRRK2 plays a fundamental role in Rab GTPase coordination of vesicular trafficking.

## Results

### Rab38 increases LRRK2’s phosphorylation of Rab10 at Thr73

Rab32 and Rab38 bind LRRK2 *in vitro* and are highly homologous to Rab29;^14^ thus, we hypothesized that Rab32 and Rab38 might regulate LRRK2 function. We expressed GFP-LRRK2 and either HA-tagged Rab29, Rab32, or Rab38 in HEK-293T cells. Autophosphorylation of LRRK2 at Ser1292 (pS1292-LRRK2) and phosphorylation of Rab10 at Thr73 (pT73-Rab10) were quantified as robust and specific measures of LRRK2 kinase activity.^11^ Overexpressed HA-Rab29 robustly increased phosphorylation of S1292-LRRK2 and T73-Rab10 by wild type (WT) or PD-mutant (R1441G or G2019S) GFP-LRRK2 (Fig. 1a, b), consistent with Purlyte et al.^11^ HA-Rab32 or HA-Rab38 did not increase pS1292-LRRK2 of WT or PD-mutant GFP-LRRK2, also consistent with Purlyte et al. Notably, both overexpressed HA-Rab32 and HA-Rab38 increased WT and PD-mutant GFP-LRRK2’s phosphorylation of T73-Rab10, with Rab32 increasing pT73-Rab10 ∼2-4 fold and HA-Rab38 increasing pT73-Rab10 ∼4-5 fold (vs. ∼4-8-fold for Rab29; Fig. 1b).

**Figure 1.**
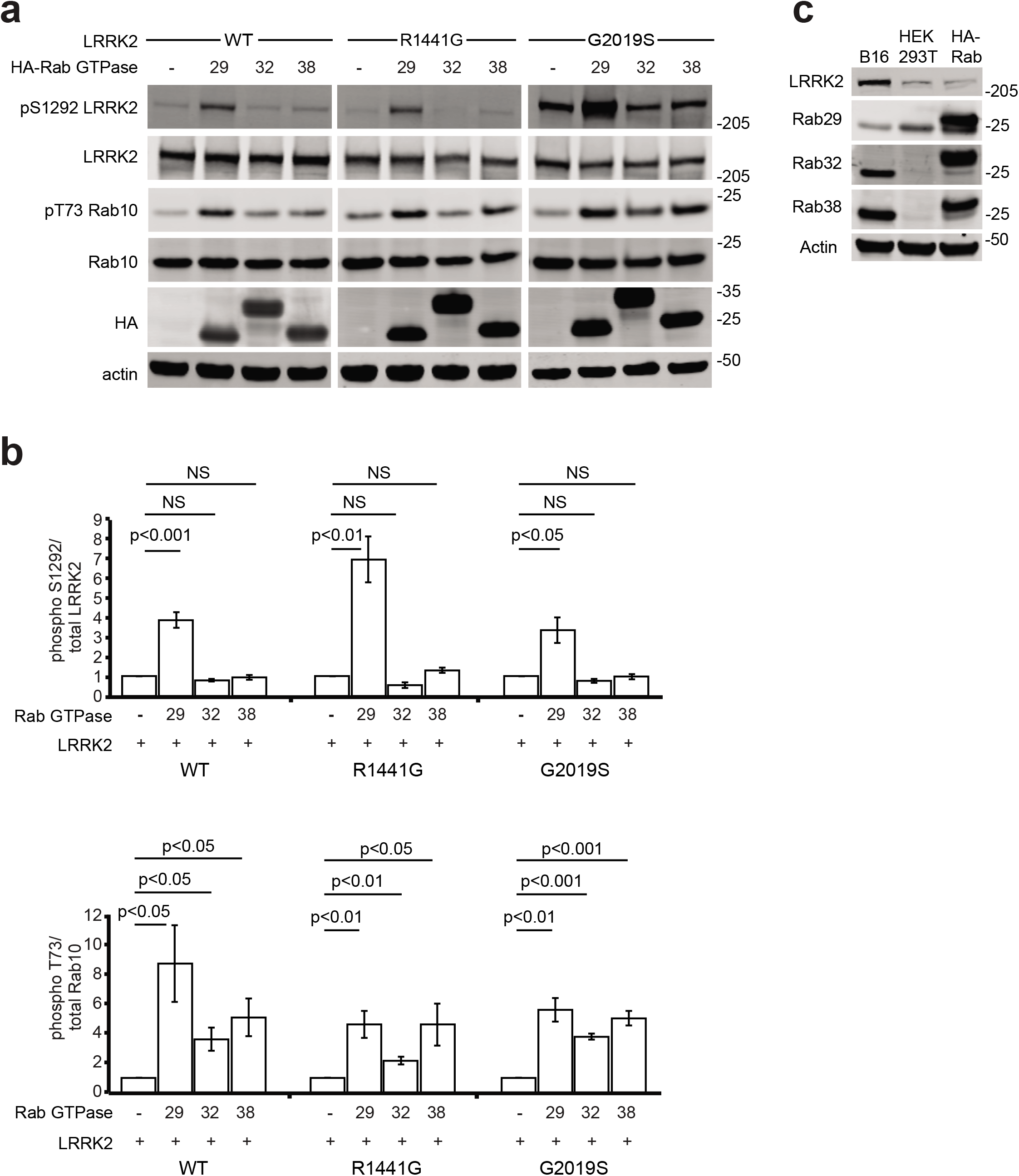
Overexpressed Rab38 and Rab32 increase LRRK2’s phosphorylation of Rab10 in HEK-293T cells. (A) Immunoblot of phosphorylated Ser1292 and Rab10 at T73 in the presence of HA-tagged Rab29, Rab32, and Rab38 for wild type LRRK2 (left), LRRK2 PD-mutant R1441G (middle), and LRRK2 PD-mutant G2019S (right) in HEK-293T cells. (B) Quantification of LRRK2 autophosphorylation and Rab10 phosphorylation in (A). (C) Immunoblot representation of endogenous Rab29, Rab32, and Rab38 levels in B16-F10 and WT HEK-293T cells relative to overexpressed HA-tagged Rab proteins in WT HEK-293T cells. Quantifications show the mean value from 4 independent replicates with error bars showing standard error of the mean. Significance testing for panel B was performed using Kruskal-Wallis test with post-hoc Dunn correction when applicable.

### Endogenous Rab38 increases LRRK2’s phosphorylation of substrate Rabs in melanocytes

HEK-293T cells do not produce appreciable Rab32 or Rab38 and express low levels of endogenous LRRK2. In our hands, RAW264.7 macrophages strongly expressed Rab29 and Rab32 but showed no expression of endogenous Rab38 by immunoblot (data not shown). In contrast, B16-F10 mouse melanoma cells (“B16 cells”) produce melanosomes (e.g. Fig. S3), indicating that pathways of lysosome-related organelle (LRO) biogenesis are active, and express high levels of endogenous Rab32, Rab38, and LRRK2 (Fig. 1c). We used siRNA to knockdown Rab29, Rab32, Rab38 or LRRK2 in B16 cells and quantified endogenous pT73-Rab10 (Fig. 2a-b, S1a). LRRK2 knockdown (“kd”) reduced pT73-Rab10 to ∼30% of scrambled control siRNA. Importantly, Rab38 knockdown also reduced pT73-Rab10 (to ∼60% of control) while Rab29 or Rab32 knockdown did not (see Fig. 2b legend for exact quantification). Rab38 is therefore the first identified Rab protein to control LRRK2 kinase function under physiologic conditions.

**Figure 2.**
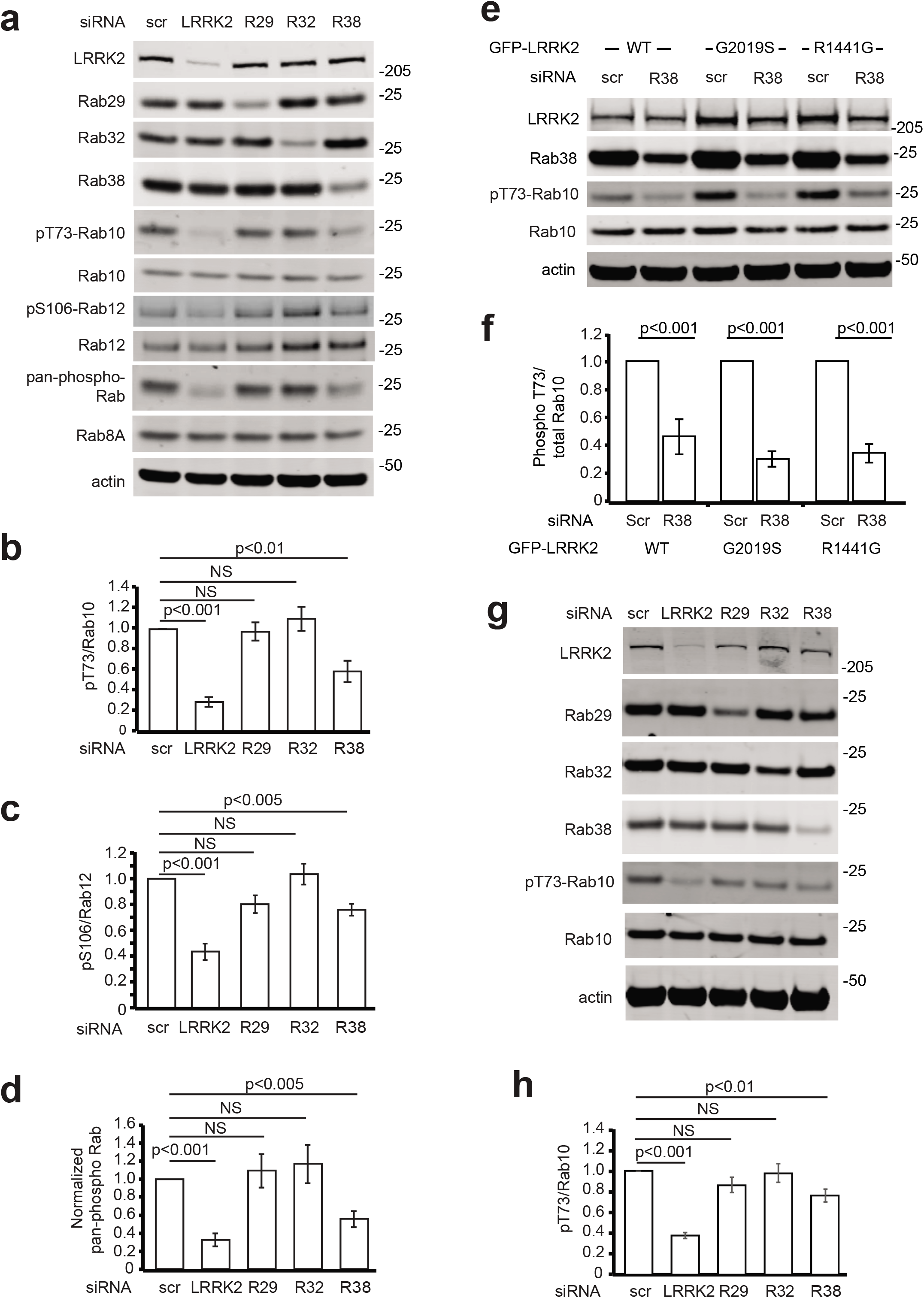
Rab38 but not Rab29 or Rab32 knockdown decreases phosphorylation of LRRK2’s substrate Rabs in melanocytes. (A) Representative immunoblot of B16 melanocytes following knockdown of LRRK2, Rab29, Rab32, and Rab38. (B) Quantification of endogenous phosphorylated Rab10 levels from 6 independent experiments. pThr73-Rab10/total Rab10 = 29% +/- 5% in LRRK2 kd, 59% +/- 11% in Rab38 kd, 97% +/- 9% in Rab29 kd, 110% +/- 12% in Rab32 kd (mean +/- SEM). Knockdown quantified in Fig. S1a. (C) Quantification of endogenous phosphorylated Rab12 levels from 4 independent experiments. pS106-Rab12/total Rab12 = 52% +/- 7% in LRRK2 kd, 84% +/- 6% in Rab38 kd, 101% +/- 14% in Rab29 kd:, and 123% +/- 13% in Rab32 kd (mean +/- SEM). Knockdown quantified in Fig. S1b. (D) Quantification of pan-phosphorylated Rab levels from 6 independent experiments using Abcam ab231706 normalized to total protein level as measured by BCA. pan-phospho Rab 32% +/- 7% in LRRK2 kd, in Rab38 kd: 56% +/- 9, in Rab29 kd: 109% +/- 19%, and in Rab32 kd: 117% +/- 21% (mean +/- SEM). Knockdown quantified in Fig. S1a. (E) Representative immunoblot of B16 melanocytes in the presence of transiently transfected GFP-LRRK2 WT (left), G2019S (middle), and R1441G (right). (F) Quantification of endogenous phosphorylated Rab10 levels in the presence of exogenous WT, G2019S, or R1441G GFP-LRRK2 from 7 independent experiments. pThr73-Rab10/total Rab10 = 46% +/- 5% in WT GFP-LRRK2, 30% +/- 2% in G2019S GFP-LRRK2, 34% +/- 2% in R1441G GFP-LRRK2 (mean +/- SEM). Knockdown quantified in Fig. S1c. (G) Representative immunoblot of melan-Ink4a cells following knockdown of LRRK2, Rab29, Rab32, and Rab38. (H) Quantification of endogenous phosphorylated Rab10 levels (pThr73-Rab10/total Rab10) from 9 independent experiments. pThr73-Rab10/total Rab10 = 64% +/- 12% in Rab38 kd, 105% +/- 11% in Rab29 kd, 115% +/- 12% in Rab32 kd (mean +/- SEM). Knockdown quantified in Fig. S1d. All quantifications show mean with error bars showing standard error of the mean. Significance testing for panels B-D, F, and H was performed using a two-tailed Student’s t-test and Bonferroni correction for multiple comparisons.

To test if Rab38 knockdown in B16 cells decreases phosphorylation of LRRK2 substrate Rabs more generally, we measured Rab phosphorylation using additional LRRK2 phosphosite-specific antibodies including phospho-Ser106 Rab12 (pS106-Rab12) and phospho-Thr72 Rab8A (pT72-Rab8A, Abcam ab231706). Abcam ab231706, although developed using a pT72-Rab8A antigen, is not specific to this phosphorylation site and reacts with LRRK2 phosphorylation sites on multiple Rabs, including Rab8A, Rab8B, Rab3A, Rab10, Rab35 and Rab43.^6^ We thus utilized Abcam ab231706 as a pan-phospho-Rab antibody and evaluated changes in this signal relative to total protein (from here, data with this antibody is presented as “pan-phospho-Rab”). Both pS106-Rab12 (Fig. 2c, Fig S1b) and pan-phospho-Rab (Fig. 2d, Fig S1a) showed similar results to pThr73-Rab10, with LRRK2 or Rab38 knockdown but not Rab29 or Rab32 knockdown decreasing pS106-Rab12 and pan-phospho-Rab signal relative to scrambled control siRNA (see Fig. 2c, d legend for exact quantification).

To test the effect of Rab38 on PD-mutant LRRK2, Rab38 knockdown was performed in B16 cells expressing exogenous WT, G2019S, or R1441G GFP-LRRK2 (Fig. 2e). Rab38 knockdown decreased phosphorylation of T73-Rab10 by overexpressed WT, G2019S, and R1441G GFP-LRRK2 (Fig. 2f, Fig. S1c). Finally, we tested the effect of Rab38 knockdown on LRRK2 kinase activity in the benign melanocytic cell line, melan-Ink4a, which is diploid and syngeneic with B16-F10.^33^ In melan-Ink4a cells, pThr73-Rab10 was decreased by LRRK2 or Rab38 knockdown but not Rab29 or Rab32 knockdown (Fig. 2g,h). In summary, endogenous Rab38 but not Rab32 or Rab29 controls endogenous, overexpressed, and PD-mutant LRRK2’s phosphorylation of substrate Rab proteins in melanocytic cells.

### Endogenous Rab38 drives LRRK2 membrane association

In non-melanocytic HEK-293T and HeLa cells, ∼10% of LRRK2 is membrane-associated and the majority is cytoplasmic.^34, 35^ In these cells, Rab29 overexpression recruits overexpressed LRRK2 to Golgi and pericentriolar membranes, activating LRRK2’s kinase and causing LRRK2 kinase-dependent pericentriolar accumulation of pT73-Rab10.^7, 11, 36^ Because endogenous Rab38 can increase LRRK2’s phosphorylation of Rab substrates, we hypothesized that endogenous Rab38 might drive LRRK2’s membrane association. We therefore isolated membrane and cytoplasmic fractions of B16 cells following Rab38 or control siRNA knockdown (Fig. 3a). Using chemiluminescence to visualize the low levels of endogenous LRRK2, we observed that the majority of endogenous LRRK2 in B16 cells was cytoplasmic and a smaller portion was membrane-associated. Upon Rab38 knockdown, the fraction of LRRK2 in the cytoplasm reproducibly increased while the membrane-associated fraction decreased (Fig. 3a, blot representative of 4 replicates).

**Figure 3.**
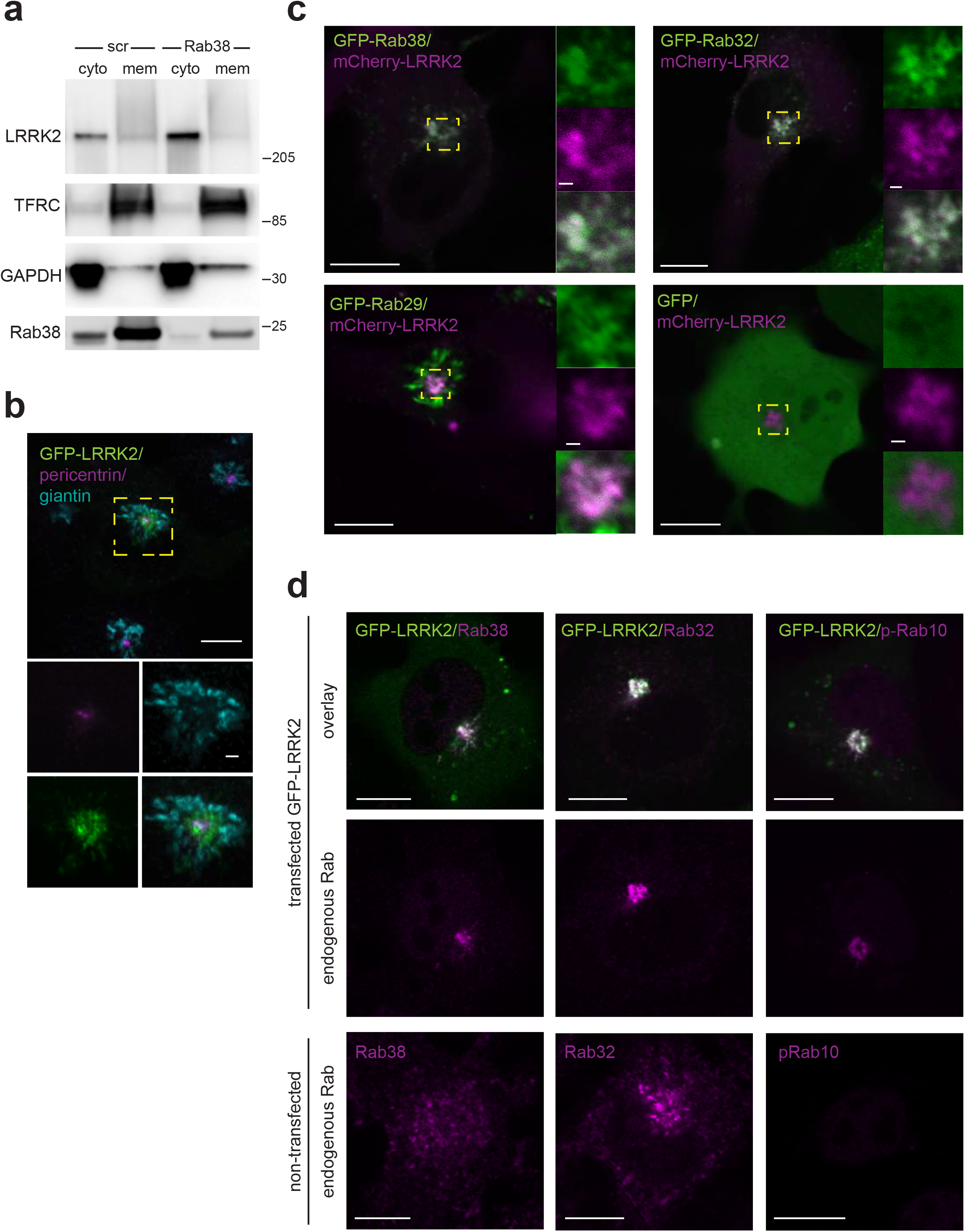
Overexpressed LRRK2 accumulates in a pericentriolar location with endogenous Rab38, Rab32, and phosphorylated Rab10. (A) Chemiluminescent immunoblot of endogenous LRRK2 in cytoplasmic and membrane fractions of B16 cells with scrambled siRNA or endogenous Rab38 knocked down by targeted siRNA. Result is representative of 4 replicates. (B) Immunofluorescence confocal microscopy of GFP-LRRK2 clustered at pericentriolar punctae in B16 cells. Insets show higher magnification and isolated channels of the region identified by the yellow box. LRRK2 is shown in green, pericentrin in magenta, and giantin in blue. (C) Live-cell confocal microscopy of mCherry-LRRK2 (magenta) and GFP-tagged Rab proteins (green) in B16 cells (overlay is white). Top panels show GFP-Rab32 (left) and GFP-Rab38 (right) accumulating at pericentriolar region with mCherry-LRRK2. Bottom panels show GFP-Rab29 (left) is Golgi-localized and GFP alone (right) is cytoplasmic in the presence of mCherry-LRRK2. Insets show higher magnification of region in yellow box. (D) Immunofluorescence confocal microscopy of endogenous Rab32, Rab38, and phospho-T73 Rab10 with and without GFP-LRRK2 in B16 cells. Top row: overlay of GFP-LRRK2 (green) and each Rab (purple) at pericentriolar region. Middle row: isolated endogenous Rab protein channels corresponding to images in top row. Bottom row: Cellular distribution of Rab38, Rab32, and phospho-Rab10 in absence of GFP-LRRK2. Scale bars = 10 µm.

### Rab38 regulates pericentriolar recruitment of GFP-LRRK2 in B16 melanocytes

We wished to evaluate subcellular localization of endogenous LRRK2 in melanocytes using immunofluorescence (IF); however, no antibodies identified endogenous LRRK2 by IF in B16 cells (not shown). Thus far, endogenous LRRK2 has been conclusively visualized by IF only in macrophages in which interferon-γ is utilized to augment LRRK2 expression.^3^ For our studies in melanocytes, we therefore expressed exogenous LRRK2 and restricted all microscopy to cells expressing visible but low levels of LRRK2. We note that in B16 cells, highly overexpressed LRRK2 sometimes formed aggregates and skein-like structures of unclear physiologic relevance, identical to its behavior in other cell lines.^11^

In non-melanocytic cells, GFP-LRRK2’s cytoplasmic localization is so prominent that cytosolic depletion is required to visualize membrane-bound LRRK2.^15^ In contrast, in B16 cells, GFP-LRRK2 formed clustered perinuclear punctae visible without cytosolic depletion (Fig. 3b). IF for pericentrin (centrosomal marker) and giantin (Golgi marker) revealed that these punctae are pericentriolar. mCherry-LRRK2 expression recruited GFP-Rab38 and GFP-Rab32 to the pericentriolar punctae (Fig. 3c and Fig. S2). In the absence of mCherry-LRRK2, GFP-Rab38 and GFP-Rab32 localized to melanosomes and dispersed perinuclear vesicles (Fig. S3). Consistently, endogenous Rab32, Rab38, and phosphorylated (pT73) Rab10 also accumulated at pericentriolar punctae in the presence, but not absence, of exogenous LRRK2 (Fig. 3d). Localization of GFP-Rab29 (Golgi)^11^ and GFP (cytoplasmic with some nuclear staining)^37^ did not change upon mCherry-LRRK2 expression (Fig. 3c and Fig. S3). Endogenous Rab29 was not visible using commercial Rab29 antibodies (Abcam ab256526, Santa Cruz 81924), consistent with some prior literature^11^ but in conflict to data using the non-commercial version of ab256526.^19^

Kinase activity of exogenous LRRK2 was required for pericentriolar accumulation of LRRK2 in B16 cells and subsequent recruitment of Rab32, Rab38, and pT73-Rab10: Kinase-inactive mutants GFP-LRRK2 T1348N and GFP-LRRK2 D2017A were diffusely cytoplasmic (Fig. 4a) and essentially no cells expressing LRRK2 T1348N or D2017A accumulated pericentriolar LRRK2 or Rab32 (Fig. 4b). Exogenous kinase-inactive LRRK2 also did not cause pericentriolar recruitment of endogenous Rab32, Rab38, or pT73-Rab10 (not shown). Pericentriolar accumulation of exogenous LRRK2 is therefore a robust readout reflecting LRRK2 kinase function, though we note that the physiologic relevance of this accumulation is unclear.

**Figure 4.**
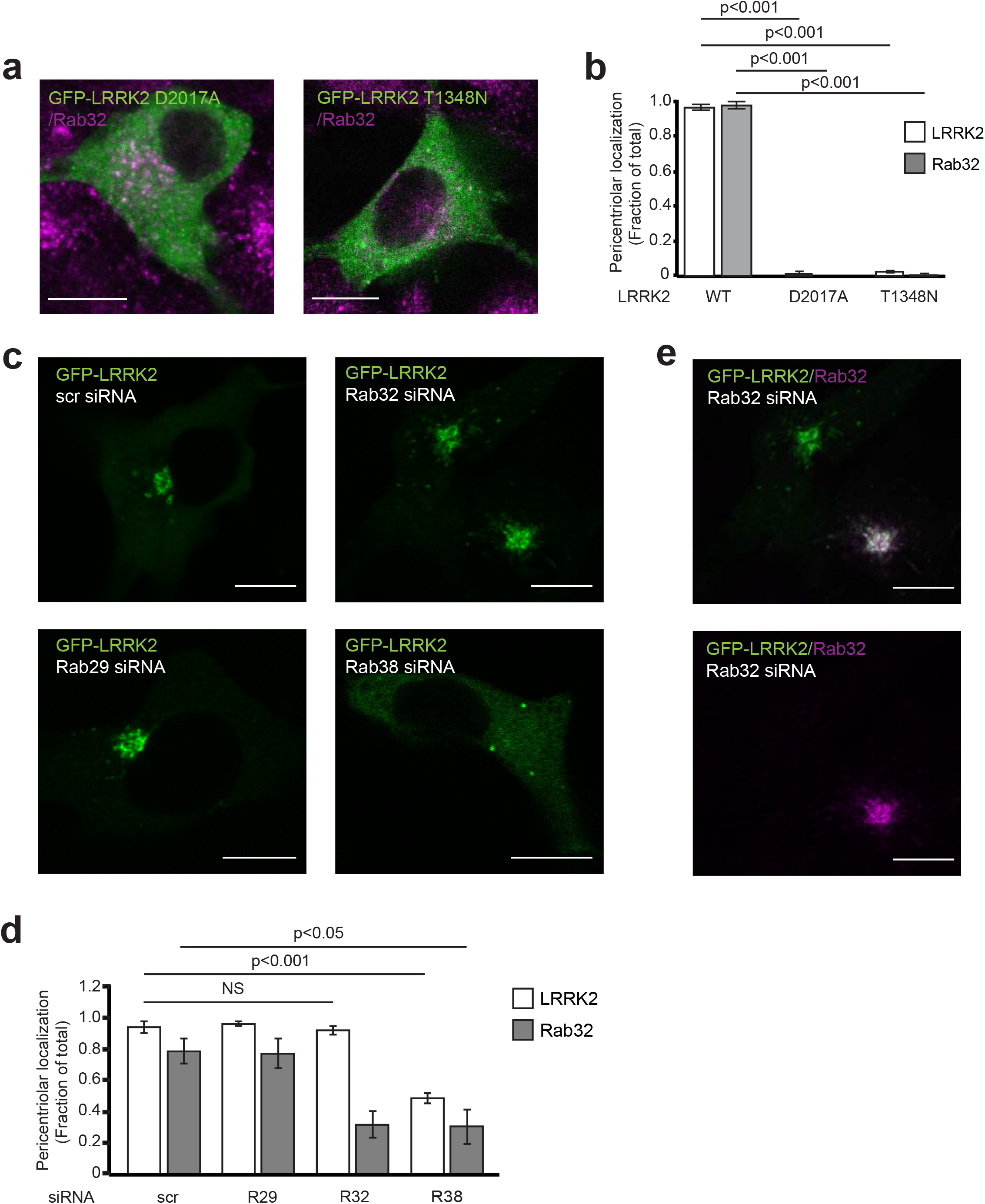
Endogenous Rab38 and enzymatically active LRRK2 drive pericentriolar recruitment of overexpressed LRRK2. (A) Immunofluorescence confocal microscopy of endogenous Rab32 (magenta) with GFP-LRRK2 D2017A (kinase-dead, left panel) or GFP-LRRK2 T1348N (GTP non-binding, right) in B16 cells (overlay to white). (B) Quantification of GFP-LRRK2 (white) and Rab32 (grey) pericentriolar recruitment in (A). Quantification includes 3 replicates of ≥50 cells per LRRK2 variant. % of cells with pericentriolar GFP-LRRK2 with WT GFP-LRRK2 = 97% +/- 2%, with T1348N GFP-LRRK2 = 3% +/- 1%, with D2017A GFP-LRRK2 = 1% +/- 1%. % of cells with pericentriolar Rab32 with WT GFP-LRRK2 = 98% +/- 2%, T1348N GFP-LRRK2 = 1% +/- 1%, D2017A GFP-LRRK2 = 0% +/- 0% (mean +/- SEM). (C) Immunofluorescence confocal microscopy of GFP-LRRK2 in B16 cells with knockdown of Rab29 (bottom left), knockdown of Rab32 (top right), knockdown of Rab38 (bottom right), or scrambled control siRNA (top left). (D) Quantification of GFP-LRRK2 (white) and Rab32 (grey) pericentriolar recruitment in (C). Quantification includes 3 replicates of ≥50 cells per knockdown condition. % of cells with pericentriolar GFP-LRRK2 was 49% +/- 3% in the presence of Rab38 knockdown, 96% +/- 1% with Rab29 knockdown, 92% +/- 3% with Rab32 knockdown, and 94% +/- 4% with control siRNA. % of cells with pericentriolar Rab32 was 31% +/- 11% in the presence of Rab38 kd, 32% +/- 8% with Rab32 kd, 77% +/- 9% with Rab29 kd, 79% +/- 8% with scr kd (mean +/- SEM). (E) Immunofluorescence confocal microscopy of GFP-LRRK2 localizing in the perinuclear region in the presence and absence of endogenous Rab32. Top: GFP-LRRK2 (green) and Rab32 (magenta) (overlay to white) for two individual B16 cells. Bottom: isolated Rab32 channel for the same two cells. All quantifications show mean with error bars showing standard error of the mean. Significance testing for panels B and D was performed using a two-tailed Student’s t-test and Bonferroni correction for multiple comparisons. Scale bars = 10 µm.

In keeping with Rab38’s apparently unique role in regulating LRRK2 kinase function and membrane association in murine melanocytes, Rab38 knockdown, but not Rab29 or Rab32 knockdown or control siRNA, robustly inhibited pericentriolar recruitment of exogenous LRRK2 (Fig. 4c,d, Fig. S1e). We also visualized and quantified endogenous Rab32 localization after Rab38, Rab32, Rab29 or control knockdown (Fig 4d). GFP-LRRK2 remained pericentriolar in the absence of Rab32 pericentriolar accumulation (Fig. 4e, top left cell), demonstrating that Rab32 was not required. However, both exogenous LRRK2 and endogenous Rab38 were necessary for endogenous Rab32 accumulation, since Rab32 accumulation significantly decreased in the Rab38 knockdown (Fig. 4d).

### Disrupting LRRK2-Rab38 interactions decreases LRRK2 membrane association, pericentriolar recruitment, and phosphorylation of Rab10

LRRK2’s ARM domain (residues 1-660) binds Rab38 family members;^38^ thus, we predicted that LRRK2_660-2527_ should show decreased Rab38-mediated phenotypes (i.e. membrane association, pericentriolar recruitment, and phosphorylation of substrate Rabs) compared to full-length LRRK2. First, we examined the role of LRRK2_1-660_ in Rab38-mediated membrane association. After Rab38 or control siRNA knockdown, the proportion of GFP-LRRK2 in membrane vs. cytoplasm fraction was quantified. ∼10% of GFP-LRRK2 associated with membranes in the presence of control siRNA (Fig. 5a, b), consistent with other cell lines.^11^ Knockdown of Rab38 to ∼10% of baseline protein levels (Fig. S1f) decreased LRRK2 membrane association >2-fold. In contrast, Rab29 knockdown to ∼10% of baseline protein levels (Fig. S1h) did not change the proportion of LRRK2 in membrane vs. cytoplasmic fractions (Fig. S4a,b).

**Figure 5.**
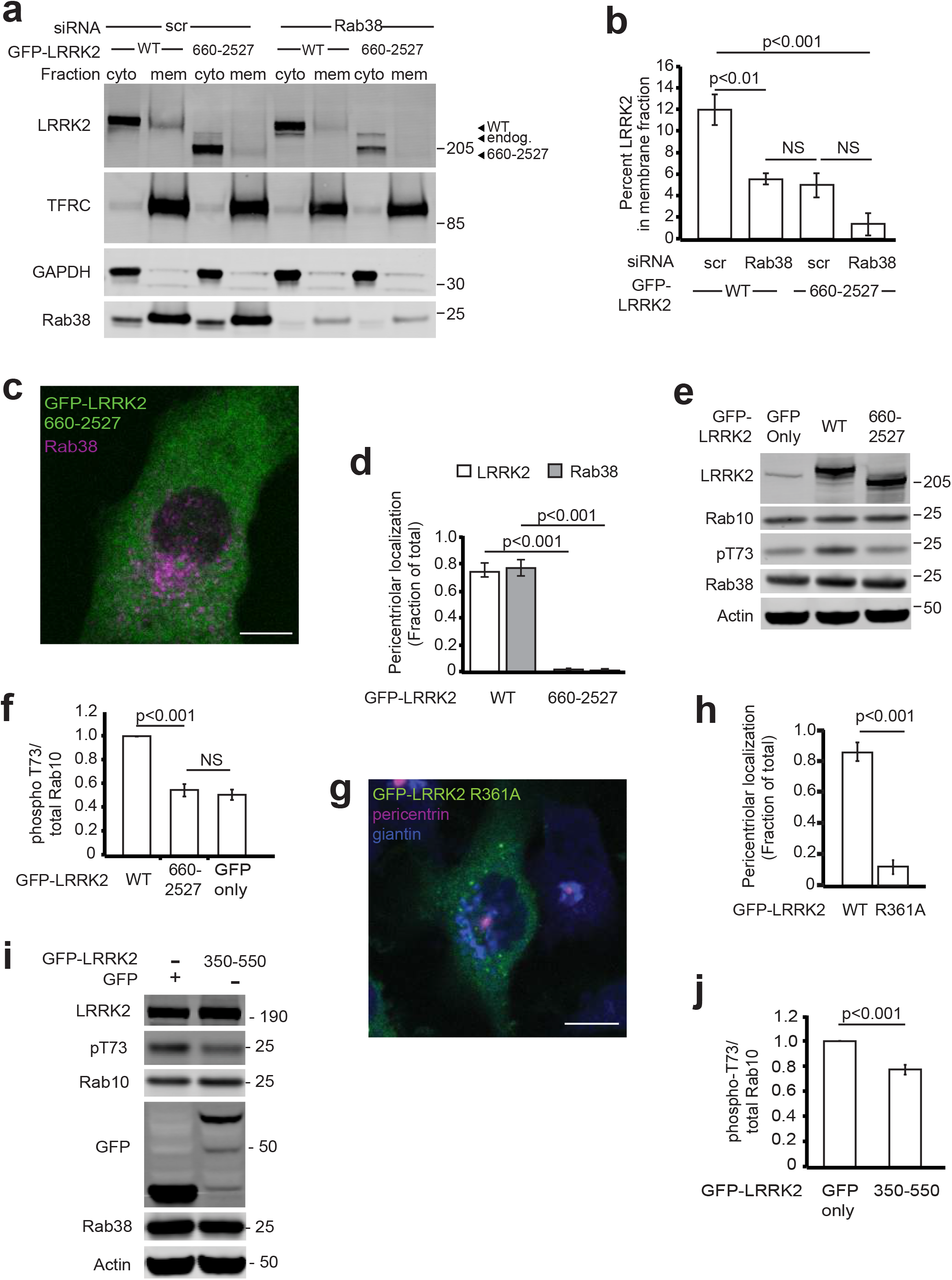
Disruption of Rab38 binding to LRRK2’s armadillo domain decreases LRRK2’s membrane association, pericentriolar recruitment, and endogenous kinase activity. (A) Immunoblot of cytoplasmic and membrane fractions of B16 cells expressing GFP-LRRK2 WT (full-length) or GFP-LRRK2_660-2527_ after Rab38 knockdown or scrambled control siRNA. (B) Quantification of membrane-associated GFP-LRRK2 versus GFP-LRRK2_660-2527_ from 4 independent experiments. After control siRNA, 11% +/- 1% of GFP-LRRK2 WT was membrane-associated while after Rab38 knockdown, 5% +/- 1% was membrane-associated (mean +/- SEM). After control siRNA, 11% +/- 1% of GFP-LRRK2 WT was membrane-associated versus 5% +/- 1% after Rab38 knockdown. After control siRNA, 4% +/- 1% GFP-LRRK2_660-2527_ was membrane associated versus 2% +/- 1% after Rab38 knockdown (mean +/- SEM), which was not significant. (C) Immunofluorescence microscopy of GFP-LRRK2_660-2527_ (green) and endogenous Rab38 (purple) in B16 cells. (D) Quantification of LRRK2 (white) and Rab38 (grey) pericentriolar recruitment in GFP-LRRK2 WT compared to GFP-LRRK2_660-2527_. Quantification includes 3 replicates of ≥50 cells per LRRK2 variant. The proportion of cells with pericentriolar GFP-LRRK2_660-2527_ was 2% +/- 1% vs. 74% +/- 5% for GFP-LRRK2 WT (mean +/- SEM). (E) Immunoblot of B16 cells expressing transiently transfected GFP alone, GFP-LRRK2 WT (full-length), and GFP-LRRK2_660-2527_; quantification of GFP-LRRK2 protein levels in Fig. S4c. (F) Quantification of endogenous phosphorylated Rab10 levels (pThr73-Rab10/total Rab10) from 4 independent experiments. pT73-Rab10 levels with GFP-LRRK2_660-2527_ = 54% +/- 5% vs GFP alone = 51% +/- 5%, with pT73-Rab10 levels in presence of GFP-LRRK2 WT set to 100%. Knockdown quantified in Fig. S4b. (G) Immunofluorescence confocal microscopy of GFP-LRRK2_R361A_ (green) and endogenous Rab38 (purple) in B16 cells. (H) Quantification of LRRK2 pericentriolar recruitment in GFP-LRRK2 WT versus GFP-LRRK2_R361A_. Quantification includes 3 replicates of ≥50 cells per LRRK2 variant. % of cells with pericentriolar GFP-LRRK2 R361A = 12% +/- 4% vs 85% +/- 6% for GFP-LRRK2 WT (mean +/- SEM). (I) Immunoblot of B16 cells expressing GFP alone versus GFP-LRRK2_330-550_. (J) Quantification of endogenous Rab10 phosphorylation in B16 cells expressing GFP-LRRK2 WT versus GFP-LRRK2_R361A_ from 4 independent experiments. pT73-Rab10/total Rab10 = 77% +/- 4% in presence of GFP-LRRK2_350-550_ compared to GFP alone. All quantifications show mean with error bars showing standard error of the mean. Significance testing for panels B, D, F, H, and J was performed using a two-tailed Student’s t-test and Bonferroni correction for multiple comparisons. Scale bars = 10 µm.

As predicted, the proportion of GFP-LRRK2_660-2527_ in the membrane fraction was decreased compared to full-length GFP-LRRK2. The fraction of membrane-associated GFP-LRRK2_660-2527_ did not differ in the presence vs. absence of Rab38 knockdown or compared to GFP-LRRK2 with Rab38 knockdown. These results support a model in which the Rab38-LRRK2_1-660_ interaction drives LRRK2 membrane association in B16 cells. Consistently, GFP-LRRK2_660-2527_ did not show pericentriolar recruitment when expressed in B16 cells (Fig 5c, d). Expression of GFP-LRRK2_660-2527_ also failed to augment phosphorylation of endogenous Rab10 relative to GFP alone (Fig. 5e, f; Fig. S4c).

In vitro studies using purified proteins demonstrate that LRRK2_386-392_ is particularly important for the LRRK2-Rab38 interaction.^14^ Similar to the very recent work defining amino acids mediating the Rab29-LRRK2 interaction,^15^ we used ColabFold^39^ AlphaFold2^40^ to model LRRK2-Rab38 interacting domains to pinpoint amino acids most important for this interaction. Our modelling suggested that LRRK2 Arg361 can form a salt bridge with Rab38 Glu70 or Glu82 (depending on the model); therefore, the Arg361Ala point mutation should disrupt the Rab38-LRRK2 interaction (Fig.S4d). Consistently, GFP-LRRK2 R361A was diffusely cytoplasmic rather than recruited to the pericentriolar region (Fig. 5g, h). Finally, we postulated that overexpression of GFP-LRRK2_350-550_, a stable fragment encompassing the Rab38-interacting region, should inhibit endogenous LRRK2’s phosphorylation of Rab10. Overexpression of GFP-LRRK2_350-550_ decreased pT73-Rab10 levels relative to expression of GFP alone (Fig. 5i, j). In sum, these findings provide evidence that LRRK2_350-550_ is critical for Rab38 binding in vivo.

### Loss of the BLOC-3 GEF decreases LRRK2 activity

GTP binding is required for formation of a LRRK2-Rab38 complex in vitro.^14^ The guanine nucleotide exchange factor (GEF) BLOC-3 exchanges Rab-bound GDP to GTP and maintains Rab32 and Rab38 in the active, membrane-bound state.^41^ BLOC-3 is composed of two protein subunits, Hps1 and Hps4. Loss of expression of either Hps1 or Hps4 causes instability and degradation of BLOC-3.^42, 43^ We therefore predicted that knockdown of Hps1 or Hps4 in B16 cells should decrease Rab38-dependent LRRK2 cellular localization and kinase activity. Consistently, GFP-LRRK2 was diffusely cytoplasmic rather than pericentriolar in the presence of Hps1 or Hps4 knockdown (Fig. 6a,b, Fig. S1e). We additionally examined the effect of BLOC-3 deficiency on LRRK2 activity in benign melanocytes. Melan-le melanocytes contain an Hps4 Q50stop mutation that ablates BLOC-3 protein expression. Melan-le melanocytes are otherwise isogenic to wild type melan-Ink4a melanocytes.^42^ We therefore quantified LRRK2-mediated phosphorylation of Rabs in melan-le vs. melan-Ink4a cells. Phospho-T73 Rab10, phospho-S106 Rab12, and phospho-pan Rab were strikingly decreased in melan-le vs. melan-Ink4a cells (Fig. 6c, d). Levels of LRRK2 and Rab32 were not significantly different between these two cell lines though levels of Rab38 were significantly elevated in melan-le cells compared to wild type (Fig. S4d). Thus, loss of the BLOC-3 GEF, which keeps Rab38 membrane-bound and active, decreases LRRK2 kinase activity as predicted.

**Figure 6.**
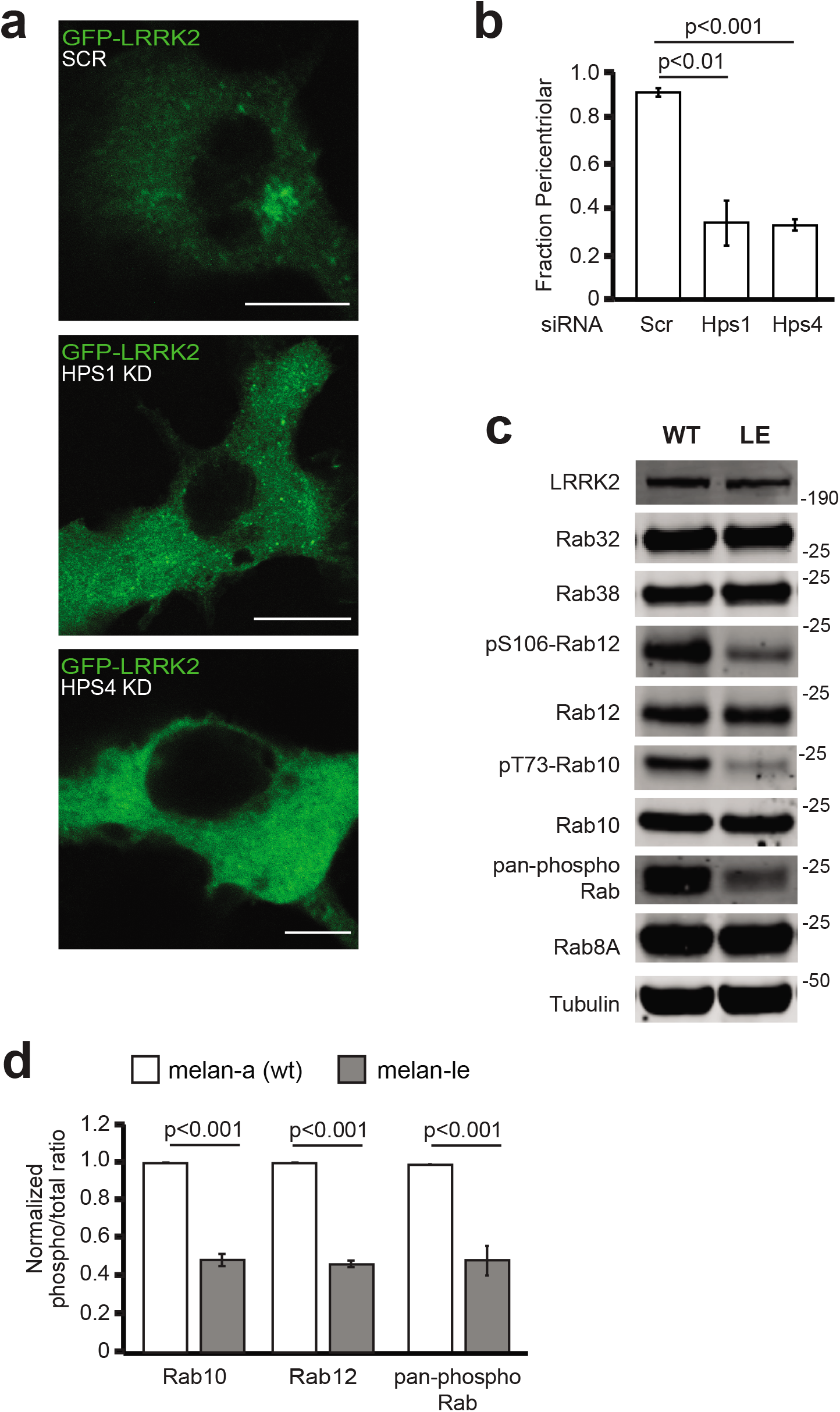
The guanine nucleotide exchange factor BLOC-3 regulates LRRK2 activity. (A) Immunofluorescence confocal microscopy of GFP-LRRK2 in B16 cells with knockdown of HPS1 (middle), HPS4 (bottom), or scrambled control siRNA (top). (B) Quantification of GFP-LRRK2 in (A). Quantification includes 3 replicates of ≥50 cells per LRRK2 variant. % of cells with pericentriolar LRRK2 with Hps1 kd = 34% +/- 10%, with Hps4 kd = 33% +/- 3%, and with control siRNA = 91% +/- 2% with control siRNA. (C) Representative immunoblot of melan-Ink4a and melan-le cells. (D) Quantification of endogenous phosphorylated Rab10, Rab12 and pan-phospho Rab levels from 6 independent experiments. Relative phosphorylation of Rabs in melan-le vs melan-Ink4a: pThr73-Rab10/total Rab10 = 42% +/- 6%, pS106-Rab12/total Rab12 = 47% +/- 2% (mean +/- SEM), pan-phospho Rab/total protein = 49% +/- 8%. Knockdown quantified in Fig. S4c. All quantifications show mean with error bars showing standard error of the mean. Significance testing for panels B and D was performed using a two-tailed Student’s t-test and Bonferroni correction for multiple comparisons. Scale bars = 10 µm.

### Loss of Rab38 function inhibits phosphorylation of under lysosomotropic stress

Upon treatment with lysosomotropic agents such as chloroquine, endogenous LRRK2 is recruited to enlarged lysosomes, leading to increased Rab10 phosphorylation by LRRK2.^19–21^ There is conflicting evidence about whether LRRK2’s recruitment requires Rab29: under lysosomal stress, Rab29 appears to mediate LRRK2 recruitment to enlarged lysosomes in Raw 264.7 murine macrophages and bone marrow derived macrophages^21^ but not in mouse embryonic fibroblasts or A549 lung epithelial cells.^19^ In B16 cells and melan-Ink4a cells, treatment with chloroquine, monensin, or nigericin all increased pThr73-Rab10 in a LRRK2-dependent manner (Fig.7a, c and data not shown). In B16 cells, we knocked down Rab29, Rab32 or Rab38 and treated with chloroquine for 3 hours. Rab38 but not Rab29 or Rab32 knockdown decreased chloroquine-mediated pThr73-Rab10 levels (Fig. 7a,b, Fig. S4e). Similarly, phosphorylation of Rab10 by LRRK2 was decreased in melan-le vs. melan-Ink4a cells after treatment with 4 μM nigericin for 2 hours (Fig. 7c, d). Thus, Rab38 is important for LRRK2 function under both basal conditions and chloroquine-induced lysosomal stress.

**Figure 7.**
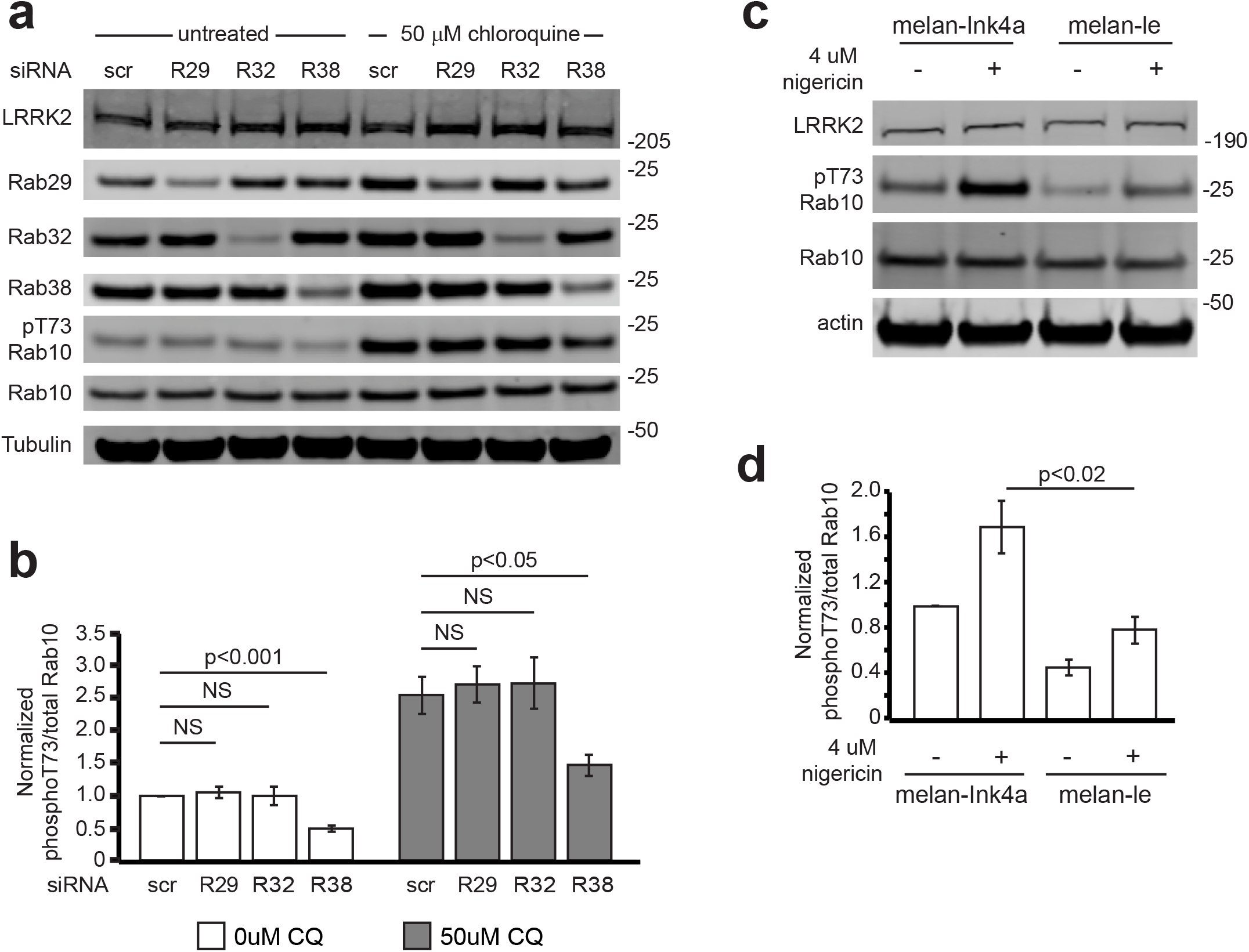
Rab38 is important in LRRK2’s response to lysosomotropic agent-mediated lysosomal stress. A) Representative immunoblot of B16 cell lysates in the presence and absence of 50 μM chloroquine treatment and with Rab29, Rab32, and Rab38 knockdown or treatment with scrambled siRNA. (B) Quantification of endogenous phosphorylated Rab10 levels (pThr73-Rab10/total Rab10) in the presence of Rab29, Rab32, and Rab38 from 6 independent experiments. Rab38 kd: 64% +/- 12, Rab29 kd: 105% +/- 11%, Rab32 kd: 115% +/- 12% (mean +/- SEM). Knockdown quantified in Fig. S4f. (C) Representative immunoblot of endogenous Rab proteins in lysate of melan-Ink4a and melan-le melanocytes in the presence or absence of 4 μM nigericin. (D) Quantification of endogenous phosphorylated Rab10 levels (pThr73-Rab10/total Rab10) in melan-Ink4a vs. melan-le melanocytes from 4 independent experiments from part (C), relative to untreated melan-Ink4a (set to 1): melan-Ink4a + nigericin: 1.70 +/- 0.23, melan-le: 0.46 +/- 0.07, melan-le + nigericin: 0.79 +/- 0.12 (mean +/- SEM). All quantifications show mean with error bars showing standard error of the mean. Significance testing for panels B and D was performed using a two-tailed Student’s t-test and Bonferroni correction for multiple comparisons.

## Discussion

Here we show that Rab38 is a new physiologic regulator of LRRK2, controlling its membrane association, subcellular localization, and phosphorylation of Rab substrates in murine melanocytes. Rab38 knockdown decreased LRRK2 membrane association in melanocytes and disrupted pericentriolar recruitment of overexpressed LRRK2 in B16-F10 melanoma cells. Additionally, Rab38 knockdown but not Rab32 or Rab29 knockdown reduced LRRK2’s phosphorylation of substrate Rabs, such as Rab10 and Rab12, under basal conditions and lysosomotropic-agent-induced lysosomal stress. Expression of LRRK2_660-2527_, which lacks the Rab38 binding domain, or the LRRK2 R361A mutation predicted to disrupt the LRRK2-Rab38 interaction, decreased LRRK2 membrane association, pericentriolar accumulation, and substrate Rab phosphorylation. Further, knockdown/mutation of the BLOC-3 GEF, which regulates Rab38 function, inhibited LRRK2 pericentriolar accumulation and substrate Rab phosphorylation. Importantly, these studies support a physiologic role for the Rab38-LRRK2 interaction and provide the first in vivo evidence that an upstream Rab regulates endogenous LRRK2 kinase function under basal cellular conditions (summarized in Fig. 8).

**Figure 8.**
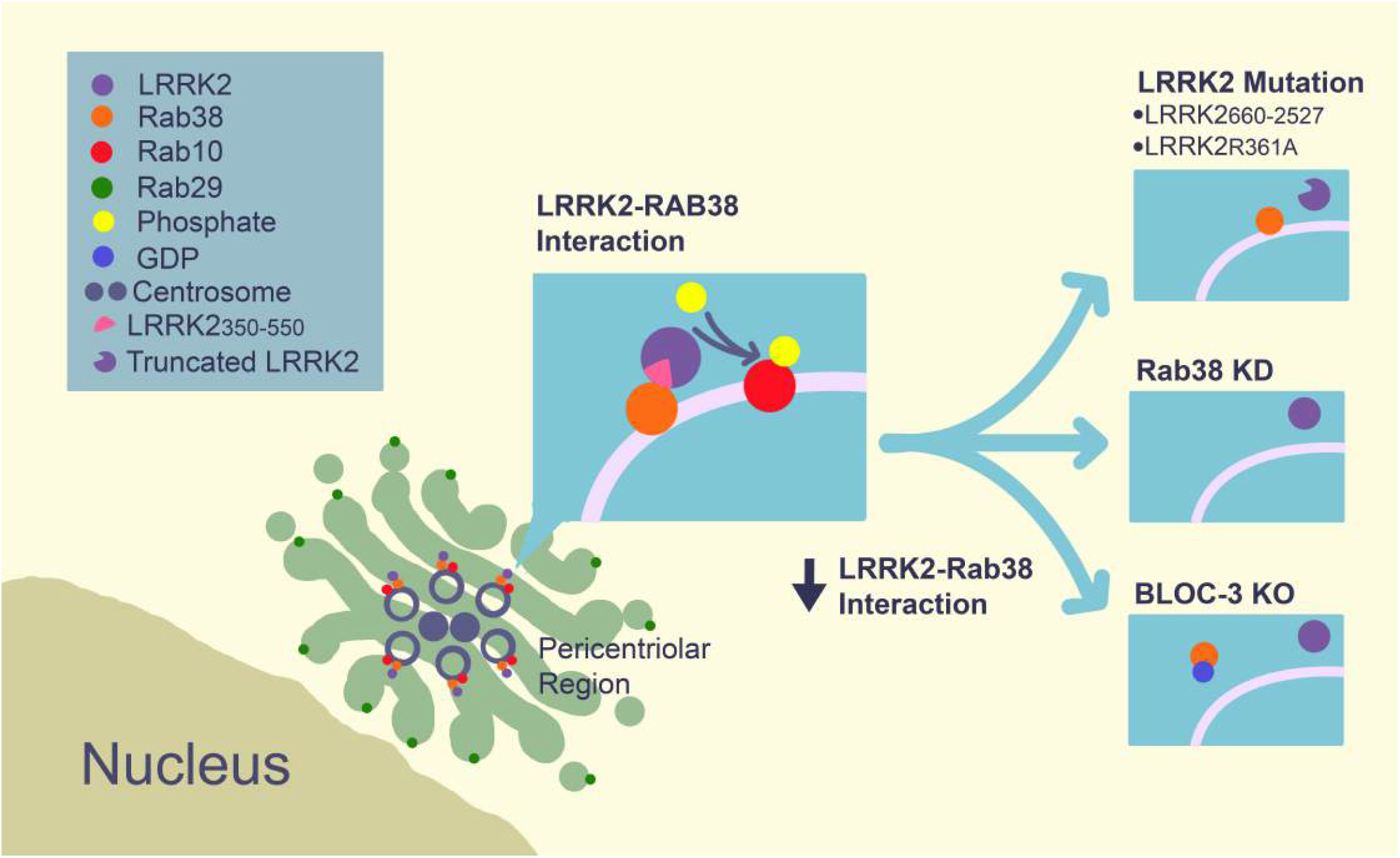
Schematic of LRRK-Rab38 interactions and functional effects.

In B16 melanocytes, we observed Rab38-dependent pericentriolar recruitment of exogenous kinase-active LRRK2 which led to pericentriolar accumulation of endogenous Rab32, Rab38 and pT73-Rab10. Overexpression of kinase-active LRRK2 was necessary for this phenotype, since endogenous Rab38, Rab32, and pT73-Rab10 did not accumulate in cells expressing only endogenous LRRK2 or with overexpression of kinase-inactive LRRK2 variants. Due to the requirement for exogenous LRRK2, we did not investigate mechanisms causing this phenotype but instead used it to validate endogenous Rab38’s regulation of LRRK2. However, we were curious about the mechanism underlying LRRK2 pericentriolar recruitment in melanocytes and what this might tell us about endogenous LRRK2 cellular function.

Interestingly, while our paper was in preparation, Vides et al. showed that LRRK2’s N-terminal lysines (in particular K18) bind specifically and with high affinity to LRRK2-phosphorylated Rab10 and Rab8 in vitro and in cells.^15^ This causes a feed-forward mechanism whereby active LRRK2 remains membrane-associated due to the interaction of its extreme N-terminus (named “Site #2”) with previously phosphorylated Rab substrates.^15^ Because Rab GTPases cluster in membrane microdomains,^44^ these authors posit that association of LRRK2 with pRab10/pRab8 via Site #2 increases the probability that LRRK2_360-450_ (“Site #1”) will interact with unphosphorylated Rab substrates. This would drive a positive feedback loop in which the effective concentration of LRRK2 molecules adjacent to Rabs would be increased, potentially allowing LRRK2 to sequentially phosphorylate multiple Rab substrates. This mechanism is entirely consistent with our observation of pericentriolar accumulation of Rab38/32/pT73-Rab10 in the setting of exogenous kinase-active but not kinase-inactive LRRK2. Although pericentriolar accumulation in B16 cells relies on exogenous LRRK2, LRRK2’s known effects on both centriolar and cilia-based phenotypes^6–9, 36^ may suggest that Rab38-mediated LRRK2 pericentriolar recruitment has physiologic relevance.

Recently, some apparently contradictory evidence regarding LRRK2 kinase activation by Rab proteins was published. In breakthrough work, the Alessi group identified LRRK2’s Rab substrates^4, 5^ and later found that overexpressed Rab29 activates LRRK2’s kinase, including increasing LRRK2 autophosphorylation of Ser1292.^11^ Their detailed follow-up studies of Rab29 knockout and overexpression mouse models; however, showed no effect of Rab29 knockout or overexpression on endogenous LRRK2 function in murine brain, lung, kidney, or spleen.^19^ They also identified no requirement for Rab29 in basal or lysosomotropic agent-augmented Rab10 phosphorylation by LRRK2 in MEFs or lung epithelial A549 cells.^19^ In contrast, the Iwatsubo group showed Rab29 to be an upstream adaptor of LRRK2 lysosomal translocation in Raw264.7 macrophages and bone-marrow derived macrophages,^21^ cells which express relatively high levels of Rab29. Additionally, detailed in vitro studies demonstrate that Rab29, Rab32, and Rab38 all interact with LRRK2’s armadillo domain with in vitro affinities between 1-3μM.^14, 15^

Here we find that, in murine melanocytes under basal and lysosomal stress conditions, Rab38 but not Rab32 or Rab29 activates LRRK2’s phosphorylation of Rab substrates and changes LRRK2 subcellular localization and membrane association. Our work adds another nuance to the already complex data about how LRRK2 membrane association is regulated. In our murine melanocyte model, Rab38 rather than Rab29 or Rab32 preferentially activates LRRK2’s ability to membrane-associate and phosphorylate Rab10. This occurs under basal as well as lysosomal stress conditions. We suggest that the most parsimonious explanation for the above results is that Rab activation of LRRK2 is highly cell-type and possibly cell-state/signal dependent. Thus, Rab29 may not contribute to LRRK2 activation under basal conditions in most cell types, but may be important in macrophages under conditions of interferon-γ signaling and lysosomal stress. In contrast, Rab38 appears important in melanocytes under both basal and lysosomal stress conditions.

Finally, our data that Rab38 but not Rab32 regulates LRRK2 kinase function and localization in mouse melanocytes adds to a body of work defining discrete roles for these highly homologous proteins (75% identical, 87% similar in humans). Rab32 and Rab38 are expressed in cells that produce lysosome-related organelles (LROs) and are critical for LRO biogenesis.^45^ Initially, their function was thought to be so similar that many works do not distinguish between them (i.e. “Rab32/38”).^24^ Studies of Rab32 and Rab38 in melanocytes thus far show largely overlapping roles in melanosome biogenesis.^45^ However, Rab32 appears more important than Rab38 in particular aspects of melanosome biogenesis such as Tyrp2 trafficking.^46^ In other LRO pathways, Rab32 but not Rab38 is required for restriction of S. typhi growth in mouse macrophages.^47^ To our knowledge, our data are the first to define a unique role for Rab38 in melanocytes that is not shared by Rab32. Murine models with loss of function mutations in Rab38 precisely phenocopy the effects of LRRK2 kinase inhibition/knockout in alveolar Type II pneumocytes.^28, 48^ Given that LRRK2 kinase inhibitors are in clinical trials,^49^ that LRRK2 inhibitor-induced lung defects are striking,^28^ and that animal studies interpreted to show reversibility of inhibitor-induced lung defects utilized short (2 week) inhibitor treatment,^29^ it is vitally important to define LRRK2’s role in normal alveolar function. More broadly, our work suggests that non-neuronal cell types expressing high levels of LRRK2 are tractable systems which can help define potentially disease-relevant LRRK2 cellular functions.

## Methods

### Plasmids and cloning

The following plasmids were either purchased from or generously donated by Dr. Dario Alessi (MRC PPU, University of Dundee, U.K.): pcDNA5 FRT/TO mCherry-LRRK2 wildtype (DU52361), pcDNA5 FRT/TO GFP-LRRK2 wildtype (DU13363), pcDNA5 FRT/TO GFP-LRRK2 L350-L550 (DU68397), pcDNA5 FRT/TO, pCMV5D-HA-Rab29 (DU50222), pCMV5D-HA-Rab32 (DU52622), pCMV5D-HA-Rab38 (DU52517, pCMV5D-GFP-Rab29 (DU50223), GFP-LRRK2 L350-L550 R361E.^15^ EGFP-Rab32 was a gift from Marci Scidmore (Addgene plasmid # 49611); DV-GFP-Rab38-WT-pDEST53 was a gift from William Pavan (Addgene plasmid # 15669).^23^ pcDNA5 FRT/TO GFP-LRRK2 660-2527 was generously donated by Dr. Jeremy Nichols and sequence verified. Full-length GFP-LRRK2 harboring an R361A mutation was generated by site-directed mutagenesis of pcDNA5 FRT/TO GFP-LRRK2 (DU13363) using the QuikChange Lightning Site-Directed Mutagenesis kit (Agilent Technologies, 210518) according to the manufacturer’s protocol and the following mutagenic primers, then sequence verified:

5’-CAAAGCATTAACGTGGCATGCCAAGAACAAGCACGTGCAGG -3’

5’-CCTGCACGTGCTTGTTCTTGGCATGCCACGTTAATGCTTTG -3’

### Cell culture, transfection, and treatments

All cell lines were grown at 37°C in a humidified atmosphere with 5% CO_2_ except melan-Ink4a, and melan-le cells (“light ear”, *Hps4^-/-^)*, which were grown at 10% CO_2_. Doxycycline-inducible GFP-LRRK2 HEK293T cell lines were cultured in DMEM+10% tetracycline-free FBS (Biowest, S1620 for all lines) containing 10 μg/mL blasticidin S (RPI, B12150) and 100 μg/mL hygromycin B (Gibco, 10687010). B16-F10 cells were cultured in RPMI+10% FBS. Melan-Ink4a, and melan-le cells were cultured in RPMI+10% FBS + 200 nM Phorbol 12-myristate 13-acetate (TPA) (Sigma-Aldrich, P8139).

Plasmid transfections were performed using Lipofectamine 2000 (Invitrogen, 11668019) using 2 μL transfection reagent/μg plasmid DNA. Cells were transfected when 60-80% confluent and plasmid transfections were incubated for 6-24 hrs. For Rab GTPase overexpression experiments in dox-inducible GFP-LRRK2 HEK293T experiments, cells were induced to express LRRK2 with 1 μg/mL of doxycycline for 18-24 hrs.

### siRNA knockdown

Cells were transfected with siRNA using either Lipofectamine RNAiMAX (Thermo Fisher, 13378075) or electroporation via Amaxa Nucleofection. For Lipofectamine RNAiMAX siRNA transfections, cells were transfected with siRNA at a final concentration of 10 nM using 0.3 μL transfection reagent/pmol siRNA. After 6-24 hours post transfection, the transfection media was removed and replaced with media lacking transfection reagent. Cells were incubated for 48-72 hrs post transfection and were harvested either by trypsinization or mechanical scraping, washed 1x with PBS, and either processed immediately or stored at −20°C. For siRNA transfection via electroporation, 1uM siRNA was electroporated into cells using the Amaxa Nucleofector II device and Cell Line Nucleofector Kit V (Lonza Bioscience, VCA-1003). B16-F10 cells were electroporated with program P-020, and melan-Ink4a/le cells were electroporated with program U-020. siRNA transfections were carried out with the following mouse siGENOME SMARTpool siRNAs: non-targeting scrambled control (Horizon Discovery, D-001206-13-05), LRRK2 (Horizon Discovery, M-049666), Rab29 (Horizon Discovery, M-053101), Rab32 (Horizon Discovery, M-063539), Rab38 (Horizon Discovery, M-040873).

### Cell lysis and immunoblotting

Cell pellets were lysed for 30 min at 4°C with end-over-end mixing in cold lysis buffer [50 mM tris pH 7.5 | 150 mM NaCl | 1 mM EDTA | 0.5% NP-40 | 1x protease (Roche, 11836170001) and phosphatase (Roche, 04906845001) inhibitors]. Different lysis buffers were used for cell fractionation experiments (detailed in fractionation methods section). Samples were centrifuged at 13,000 rcf for 5-10 mins at 4°C, and cleared lysates were quantified using the Pierce BCA Protein Assay Kit (Thermo Scientific, 23225). Lysates were denatured and reduced using LDS sample buffer (Invitrogen, NP0007) containing 5% beta-mercaptoethanol and heated at 70-95°C for 5-10 mins.

Protein samples (30-50 μg) were electrophoresed using NuPAGE 4-12% Bis Tris (Invitrogen, NP0321) or 3-8% Tris Acetate (Invitrogen, EA0375) polyacrylamide gels. Proteins were transferred from gels onto PVDF membrane (EMD Millipore, IPFL00010) using the Genscript eBlot L1 wet transfer system (cat# L00686). For fluorescent detection of western blots, membranes were blocked using LI-COR Intercept Blocking Buffer (cat# 927-60001). For chemiluminescent detection of western blots, membranes were blocked using 5% milk+0.5% Tween-20 in TBS (50 mM Tris pH 7.5, 150 mM NaCl). Immunoblotting was carried out using the following primary antibodies: mouse anti-LRRK2 N241 [Antibodies Inc. 75-253], rabbit anti-LRRK2 C41-2 (Abcam, ab133474), rabbit anti-LRRK2 phospho S1292 (Abcam, ab203181), mouse anti-Rab10 (Sigma, SAB5300028), rabbit anti-Rab10 phospho T73 (Abcam, ab230261), mouse anti-Rab8A (Santa Cruz, 81909), rabbit anti-Rab8A phospho T72 utilized as a pan-phospho-Rab antibody (Abcam, 230260), mouse anti-Rab12 (Santa Cruz, 515613), rabbit anti-Rab12 phospho S106 (Abcam, 256487), rabbit anti-Rab38 (Cell Signaling, 14365S), mouse anti-Rab32 (Santa Cruz, 390178), rabbit anti-Rab29 (Abcam, ab256526), rabbit anti-GFP (Abcam, ab290), mouse anti-TFRC (Abcam, ab269513), rabbit anti-GAPDH (Cell Signaling, 2118S), mouse anti-HA (Sigma, H3663), rabbit anti-actin (Cell Signaling, clone 13E5, 4970), mouse anti-actin (Cell Signaling, clone 8H10D10, 3700S) and rabbit anti-tubulin (Cell Signaling, 15115S). All primary antibodies were used at 1:1,000 dilution, except for actin and tubulin antibodies, which were used at 1:2,000 to 1:5,000 dilution, and GAPDH antibody, which was used at 1:5,000 dilution, and LRRK2 phospho S1292 antibody, which was used at 1:250 dilution. Primary antibodies were incubated either at room temperature for 3 hrs or overnight at 4°C or room temperature.

For fluorescent detection of western blots, goat anti-mouse (LI-COR, 926-32210) or goat-anti-rabbit (LI-COR, 926-68071) secondary antibodies were used at 1:10,000 dilution. For chemiluminescent detection of western blots, HRP-conjugated donkey anti-mouse (Jackson ImmunoResearch, 715-035-150) or donkey anti-rabbit (Jackson ImmunoResearch, 711-035-152) secondary antibodies were used at 1:10,000 dilution. Secondary antibodies were incubated at room temperature for 30-45 min. Fluorescent detection was carried out with a Li-COR Odyssey CLx imaging system. Chemiluminescent detection was carried out with SuperSignal West Pico PLUS (Thermo Scientific, cat# 34579) and/or Femto (Thermo Scientific, cat# 34094) ECL reagent and blots were imaged using an Azure Biosystems c300 Imager. Fluorescent immunoblot quantifications were performed with Image Studio Lite software version 5.2.

### Immunofluorescence microscopy

B16-F10 cells were cultured in RPMI+10%FBS on 35mm poly-d-lysine coated, No. 0 coverslip dishes (MatTek, P35GC-0-14-C) and incubated at 37 °C at 5% CO_2_. Cells at 40% confluence were transfected with 1 ug of GFP-tagged LRRK2 plasmid using Lipofectamine 2000 (Invitrogen, 11668019) as described for 12-15 hrs. For knockdown experiments, B16-F10 cells were electroporated with siRNA using Amaxa nucleofection as described. After transfection, cells were fixed using a 4% PFA solution [0.2M sucrose | 4% paraformaldehyde solution in PBS (Thermo Fisher, J19943-K2)], washed with 0.2% BSA + PBS solution, and blocked with a 5% donkey serum solution (Jackson ImmunoResearch, 017-3000-121) Coverslips were stained with the following primary antibodies: mouse-anti Rab32 (Santa Cruz, 390178), rabbit anti-Rab38 (gift from Santiago Di Pietro),^46^ mouse anti-Rab10 (Sigma, SAB5300028), and rabbit anti-Rab10 phospho T73 (Abcam, ab230261). All primary antibodies were treated for 1-3 hours at room temperature and used at a 1:200 dilution except Rab38, which was used at a 1:1,000 dilution. Secondary antibodies were used at a 1:200 dilution for 30 minutes at room temperature in the dark and include AlexaFluor647 AffiniPure donkey anti-mouse IgG (Jackson ImmunoResearch, 715-605-151), Rhodamine RedX (RRX) AffiniPure donkey anti-mouse IgG (Jackson ImmunoResearch, 715-295-150), AlexaFluor647 AffiniPure donkey anti-rabbit IgG (Jackson ImmunoResearch, 711-605-152), Rhodamine RedX (RRX) AffiniPure donkey anti-rabbit IgG (Jackson ImmunoResearch, 711-295-152). Both primary and secondary antibodies were diluted in a solution composed of 1% BSA and 0.3% Triton X-100 in PBS. Fixed-cell imaging was preformed using the Olympus IX-80 Fluoview 1000 laser scanning confocal microscope. A 60x oil-immersion objective with NA 1.42 was used to obtain confocal images (1,024 x 1,024 pixels). Z-stack images were acquired with a step-size of 0.5-1μm. All microscopy images were processed using the Fiji software package.

### Quantification of confocal imaging

Following transient transfection with GFP-LRRK2 WT or variants, at least 50 randomly selected healthy cells expressing low but visible levels of GFP-LRRK2 per plate were photographed. Most experiments quantified all such cells on the plate. Because GFP-LRRK2 has been shown to form filaments and non-specific aggregates when highly overexpressed, we focused on only those cells with lower GFP-LRRK2 levels (i.e. closer to physiologic LRRK2 levels) and excluded all cells with GFP-LRRK2 filaments or aggregates. Cells were counted by a blinded observer and considered to have LRRK2 perinuclear localization if there was clustered punctate fluorescence brighter than the immediate surrounding and concentrated near the nucleus. In unclear cases, the cells were counted as not having perinuclear LRRK2. The proportion of perinuclear LRRK2 and Rab32 were calculated for each replicate, and the average and standard error of the mean was reported.

### Cell fractionation

B16-F10 cells were initially transfected for 6-30 hrs with plasmid expressing GFP-LRRK2 WT or GFP-LRRK2 A660-E2527 using Lipofectamine 2000 as described. Following 6-hr plasmid transfection, cells were transfected with either scrambled siRNA or siRNA targeting Rab38 transcript using lipofectamine RNAiMAX as described. 24 hr after siRNA addition, the media was removed and replaced with fresh media. Cells were allowed to grow for an additional 24 hrs (48 hr siRNA knockdown), and were then harvested. Cells were fractionated by differential detergent lysis using proprietary buffers within the Mem-PER™ Plus Membrane Protein Extraction Kit (Thermo Scientific, cat# 89842). Cell pellets were resuspended in 100 or 250 μL Permeabilization Buffer containing protease (Roche, cat# 11836170001) and phosphatase (Roche, cat# 04906845001) inhibitors and lysed for 30 min at 4°C with rotation. Samples were centrifuged at 16,000 rcf for 15 min at 4°C, and the cleared supernatant was collected as the cytosolic fraction. To minimize cytosolic carryover contamination, pellets were washed 2-3x by resuspending in 1 mL Cell Wash Solution, centrifuging the suspension, and removing residual wash buffer. The washed pellets were resuspended in 100 or 250 ul Membrane Solubilization buffer containing protease and phosphatase inhibitors and incubated for 30 min at 4°C with rotation. Samples were centrifuged at 16,000 rcf for 15 min at 4°C, and the cleared supernatant was collected as the membrane fraction. Protein content for all fractions was quantified using the Pierce™ BCA Protein Assay kit. Sample buffer was added to 1x for all fractions, and samples were heated at 70°C for 10 min before storing at −20°C. Fractions from each sample were analyzed by western blot as described above. 8-30 μg protein from the membrane fraction was loaded onto gels. Cytosolic fraction loading volumes were equal to 25% of membrane fraction loading volumes.

### Quantification and statistical analysis

General statistical analysis was performed using Excel, R, Python, or STATA. For data for which normality was assumed (designated in figure legends), significance was evaluated using either an unpaired, two-tailed Student’s t test for two-sample comparisons or ANOVA with post hoc t-test and Bonferroni correction for three or more groups. For data for which normality could not be assumed (designated in figure legends), nonparametric testing was used, either the Mann–Whitney U test for two-sample comparison or Kruskal–Wallis followed by post hoc Dunn test with Bonferroni correction for multiple comparisons. A p-value < 0.05 was considered statistically significant.

## Acknowledgements

The authors thank Dario Alessi for plasmids, Mickey Marks for the melan-le cell line and advice on melanocytic cell culture, Dot Bennett for the melan-Ink4a cell line, and Dave Coughlin and Michael Perry for statistical advice and analyses. This work was made possible with support from grants from the Alzheimer’s Association (AACSF-20-684991) and NIH NINDS (K08NS090633) to A.H.

---

**Figure S1.**
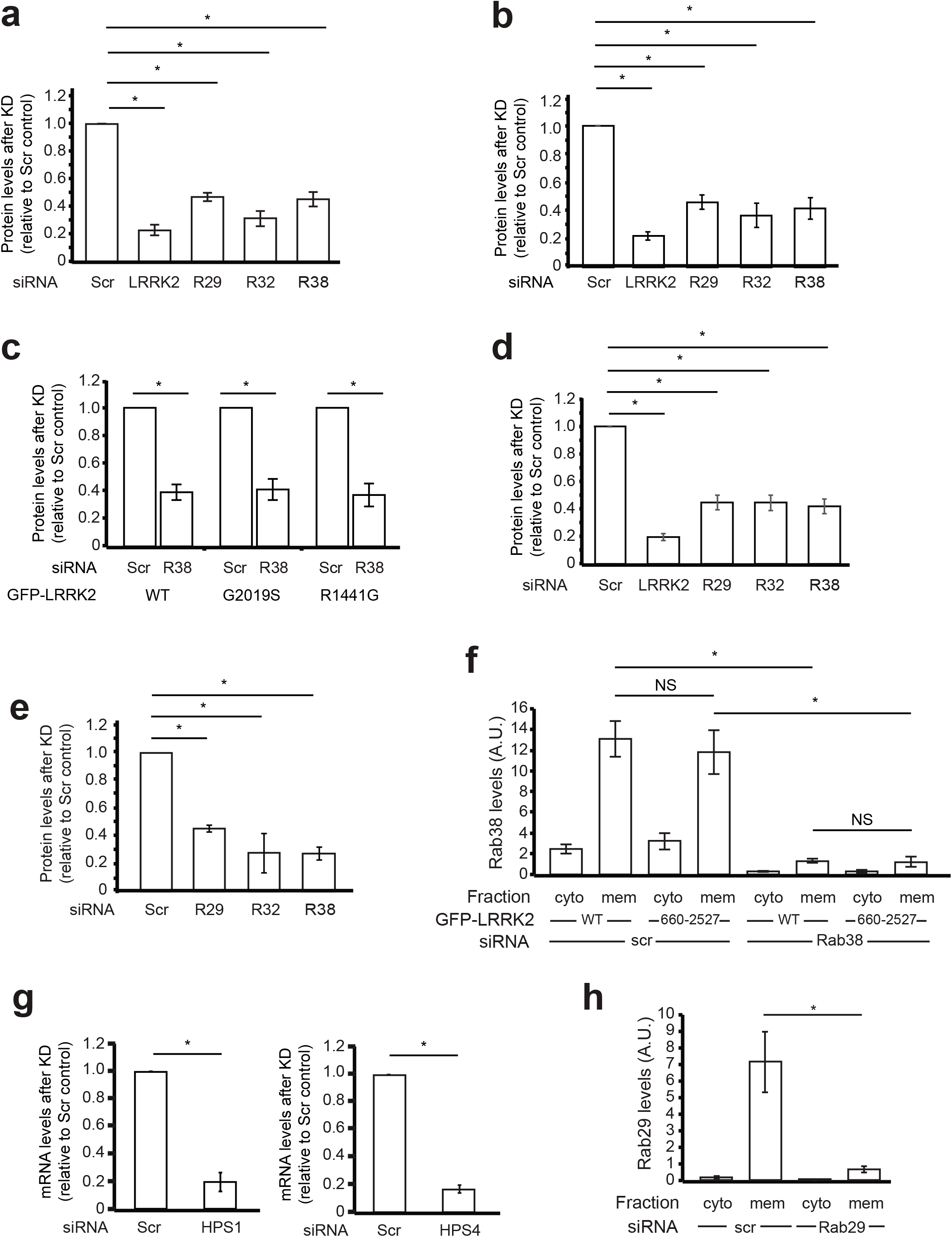
Quantification of protein or mRNA levels following siRNA knockdown. Quantification of endogenous protein (A-F) or mRNA (G) levels following siRNA knockdown of the given gene: (A, B) LRRK2, Rab29, Rab32, and Rab38 protein levels following knockdown in B16 cells with (A) showing 7 independent experiments corresponding to Fig. 2b, c and (B) showing 4 independent experiments corresponding to Fig. 2d. (C) Rab38 protein levels following knockdown in B16 cells in the presence of transient transfection of GFP-LRRK2 WT (left), G2019S (middle), and R1441G (right); 7 independent experiments corresponding to Fig. 2f. (D) LRRK2, Rab29, Rab32, and Rab38 knockdown in melan-Ink4a cells showing 8 independent knockdowns corresponding to Fig. 2h (E) Rab29, Rab32, and Rab38 levels following knockdown in B16 cells from two independent experiments corresponding to Fig. 4d. (F) Rab38 levels in cytosolic vs. membrane fractions of B16 cells following knockdown of Rab38 or scrambled control in the presence of transient transfection of GFP-LRRK2 (full-length) versus GFP-LRRK2_660-2527_; 4 independent experiments corresponding to Fig. 5b. (G) HPS1 (left) and HPS4 (right) mRNA levels following siRNA knockdown in B16 cells; 3 independent experiments corresponding to Fig. 6d. All graphs show extent of knockdown compared to scrambled control and are expressed as mean values with error bars showing standard error of the mean. Asterisks represent significant (p<0.05) t-tests results adjusted for multiple comparisons. (H) Rab29 levels in cytosolic vs. membrane fractions of B16 cells following knockdown of either Rab29 or scrambled control in the presence of transient transfection of GFP-LRRK2 (full-length); 4 independent experiments corresponding to Fig 7c.

**Figure S2.**
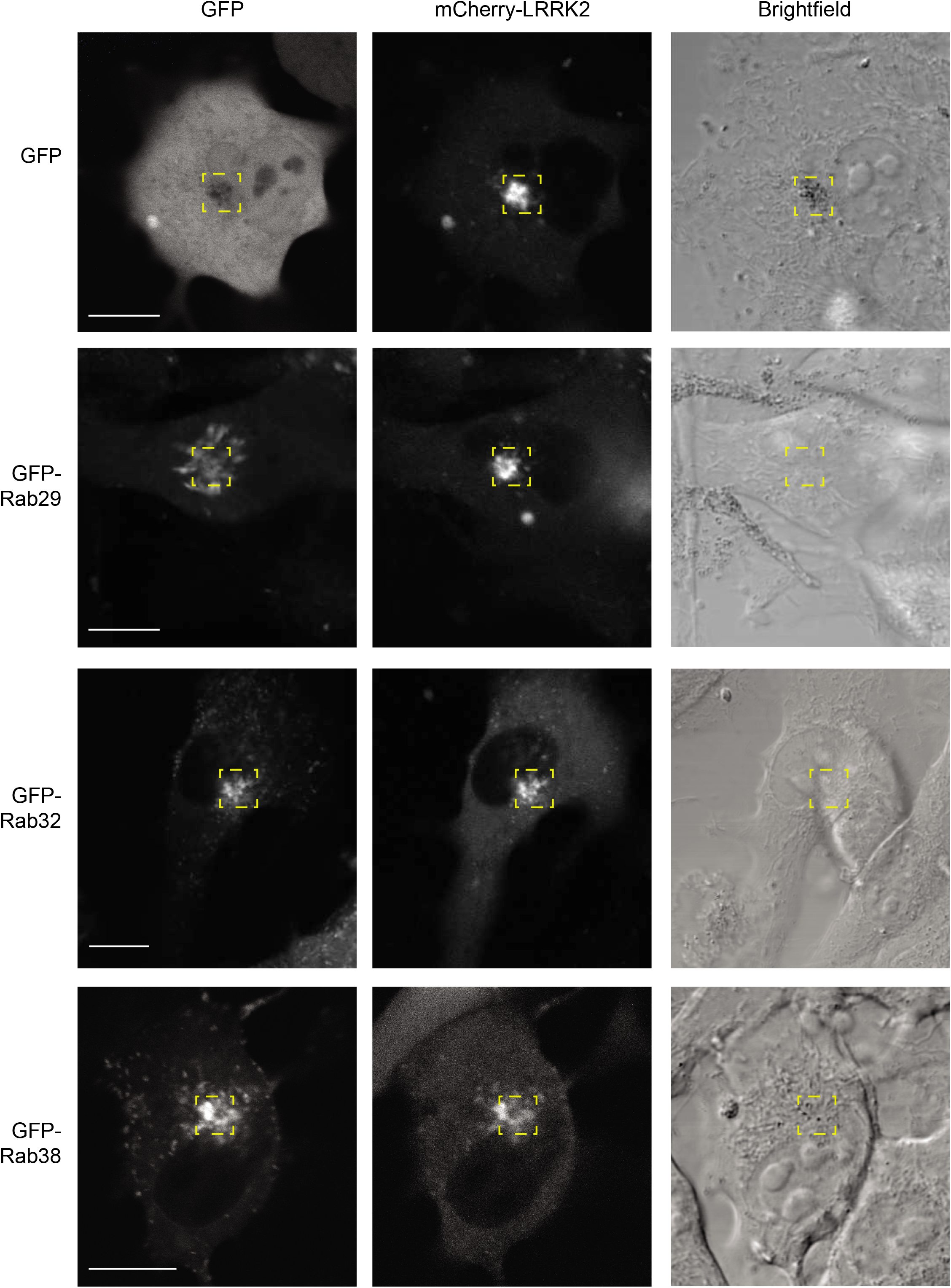
Overexpressed Rab32 and Rab38 co-localize with LRRK2 at pericentriolar region. Live-cell confocal microscopy of GFP-tagged Rab proteins (left), mCherry-LRRK2 (middle), and brightfield (right) in B16 cells. Leftmost panels show isolated GFP channels with yellow boxes highlighting GFP-tagged protein localization. Middle panels show isolated mCherry channels with yellow box highlighting mCherry-tagged LRRK2 at pericentriolar region. Rightmost panels show brightfield channel for imaged cells with yellow boxes highlighting pericentriolar area for imaged cells. Scale bars = 10 µm.

**Figure S3.**
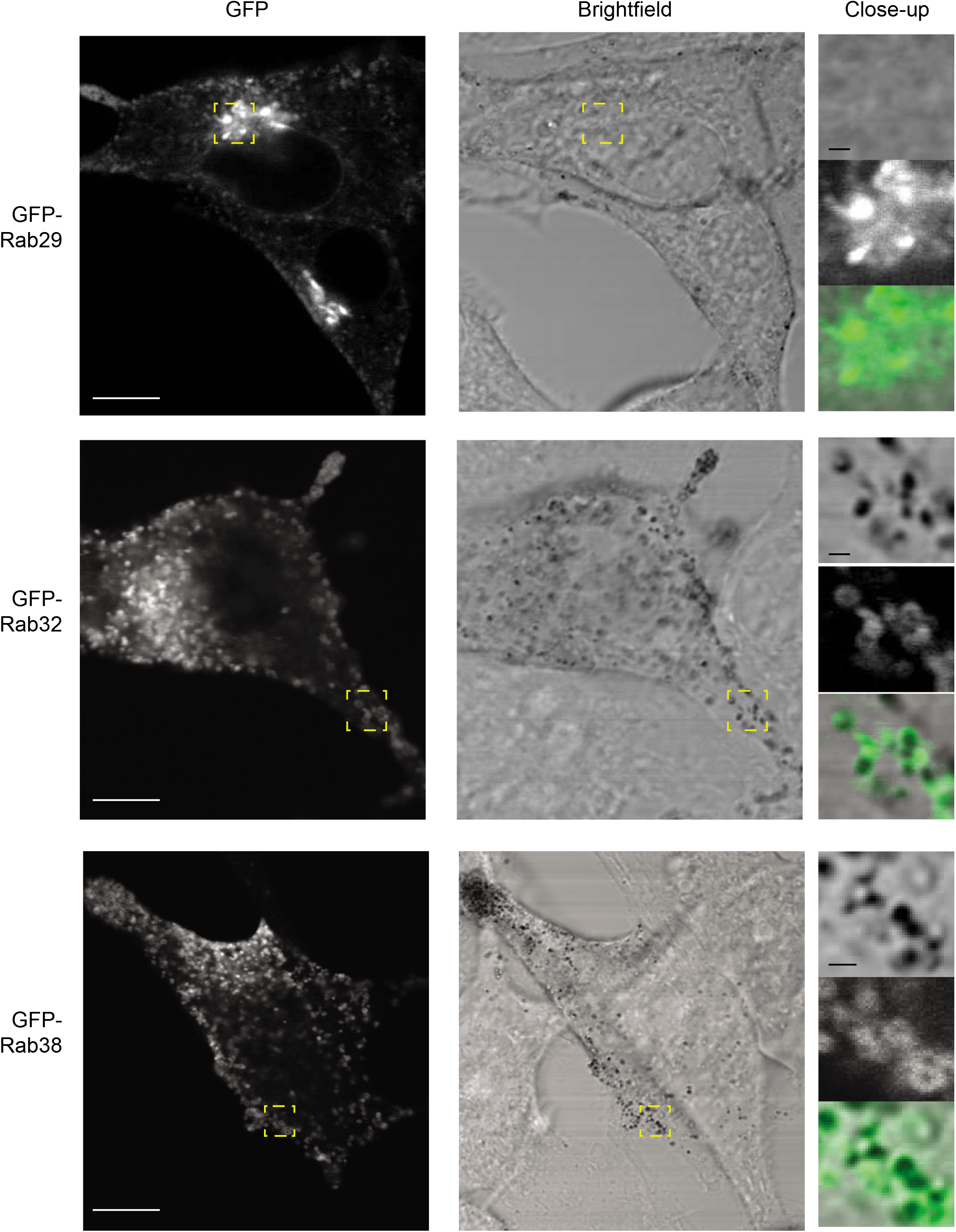
Overexpressed Rab proteins are not recruited to the pericentriolar region in the absence of overexpressed LRRK2. (A) Live-cell confocal microscopy of GFP-tagged Rab proteins in B16 cells. GFP-Rab29 (top), GFP-Rab32 (middle), GFP-Rab38 (bottom). Yellow boxes highlight area showing Rab29 at Golgi or Rab32/Rab38 co-localizing with melanosomes. Bottom panels show GFP-tagged Rab38 also at dispersed perinuclear vesicles, with yellow box Rightmost insets show close-ups of isolated brightfield (top), GFP (middle), and merged channels (bottom) for the yellow-boxed region. Scale bars = 10 µm.

**Figure S4.**
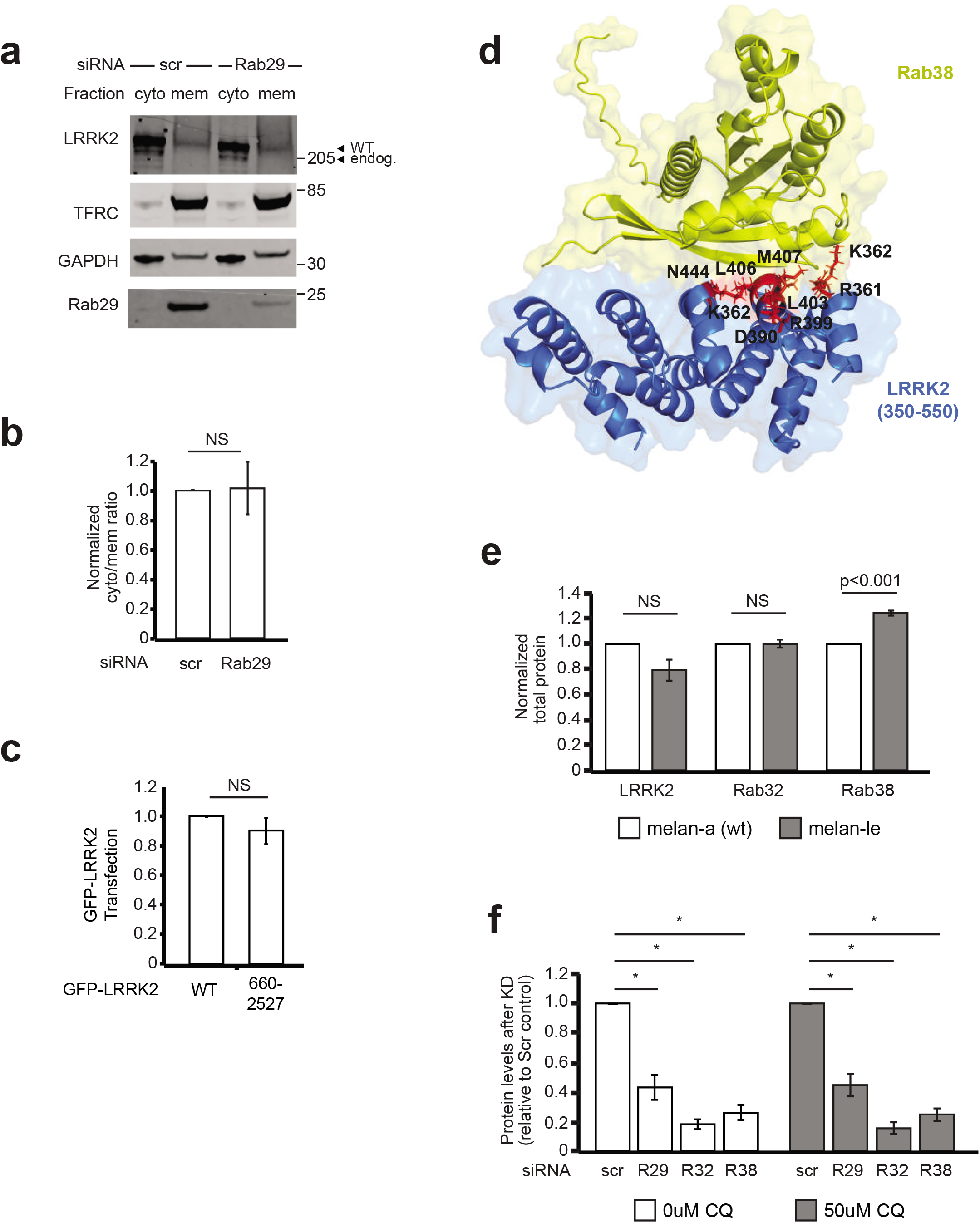
Protein levels and structural modelling of LRRK2 and Rab38. (A) Immunoblot of cytoplasmic and membrane fractions of B16 cells expressing GFP-LRRK2 WT after Rab29 knockdown or scrambled control siRNA. (B) Ratio of membrane-associated to cytoplasmic GFP-LRRK2 in Rab29 knockdown relative to scrambled siRNA control from 4 independent experiments from part A. After Rab29 knockdown, the ratio of membrane-associated to cytoplasmic GFP-LRRK2 was 1.02 +/- 0.04 (mean +/- SEM) of the scrambled control ratio (set to 1), which was not significantly different. (C) Quantification of LRRK2 protein levels for GFP-LRRK2 (full-length) versus GFP-LRRK2_660-2527_ transiently transfected into B16 cells; 7 independent experiments corresponding to Fig. 5f. (D) Model of LRRK2-Rab38 interacting domain (LRRK2_350-550_ in blue, Rab38 in yellow, predicted binding interface in red); modelling of LRRK2_350-550_ and Rab38 complex binding was performed employing ColabFold using MMseqs2 for sequence alignment, and AlphaFold2-multimer-v2 model for complex prediction; structure visualized using PyMOL. (E) LRRK2, Rab32, and Rab38 protein levels in melan-Ink4a (white) vs. melan-le (grey) quantified from the 4 independent experiments from Fig. 6b. Levels in melan-le cells relative to melan-Ink4a: LRRK2: 79% +/- 9%, Rab32: 100% +/- 2%, Rab38: 124% +/- 2% (mean +/- SEM). (F) Rab29, Rab32, and Rab38 protein levels after knockdown in non-treated (white) vs. 50uM chloroquine treated (grey) B16 cells; 4 independent experiments corresponding to Fig. 7b. Graphs show mean values with error bars showing standard error of the mean. Asterisks represent significant (p<0.05) t-tests results.

